# Comparison of phage and plasmid populations present in the gut microbiota of Parkinson’s disease patients

**DOI:** 10.1101/2023.10.23.563061

**Authors:** Alexandre Lecomte, Ilhan Cem Duru, Pia Laine, Tânia Keiko Shishido, Joni Suppula, Lars Paulin, Filip Scheperjans, Pedro Pereira, Petri Auvinen

**Affiliations:** Institute of Biotechnology, University of Helsinki, Helsinki, Finland; Department of Neurology, Helsinki University Hospital and Clinicum, University of Helsinki, Helsinki, Finland; Institute of Biotechnology, University of Helsinki, Helsinki, Finland; Department of Neurology, Helsinki University Hospital and Clinicum, University of Helsinki, Helsinki, Finland

**Author notes:** equal contributors (Pedro Pereira, Petri Auvinen).

## Abstract

The aging population worldwide is on the rise, leading to a higher number of Parkinson’s disease (PD) cases each year. PD is presently the second most prevalent neurodegenerative disease, affecting an estimated 7-10 million individuals globally. This research aimed to identify mobile genetic elements in human fecal samples using a shotgun metagenomics approach. We found over 44,000 plasmid contigs and compared plasmid populations between PD patients (n = 68) and healthy controls (n = 68). Significant associations emerged between Body Mass Index (BMI) and plasmid alpha diversity. Moreover, the gene populations present on plasmids displayed marked differences in alpha and beta diversity between PD patients and healthy controls. We identified a considerable number of phage contigs that were differentially abundant in the two groups. Moreover, we improved the continuity and identification of the protein coding regions of the phage contigs by implementing alternative genetic codes. We built a classification system based on a selection of the phages differentially abundant in the groups. A machine learning approach based on phage abundances allowed a classification of the subjects into the PD or control group with an area under curve (AUC) of 0.969.

## Introduction

Parkinson’s disease (PD) is the second most prevalent neurodegenerative disorder globally, affecting 7-10 million individuals (ref^1^ Dorsey et al., 2007). Although the underlying causes of PD remain unclear, it is widely acknowledged that the composition of the gut microbiota (GM) can affect brain health and is linked to PD (ref^2^ Scheperjans et al., 2015).

Non-motor symptoms typically manifest before the onset of characteristic motor symptoms in PD patients. One of the most prevalent non-motor symptoms is gastrointestinal dysfunction, which is frequently associated with alpha-synuclein accumulation in the enteric nervous system (ref^3^ Braak et al., 2003; ref^4^ Braak et al., 2006; ref^5^ Cersosimo et al., 2012; ref^6^ Mertsalmi et al., 2017). Progression of PD pathology to the central nervous system indicates involvement of the microbiota-gut-brain axis (ref^7^ Perez-Pardo et al., 2017; ref^8^ Wang et al., 2016).

The microbiota-gut-brain axis is a two-way communication pathway linking the GM and the central nervous system. This pathway plays a role in neural development, neuroinflammation, and modulation of complex behaviors (ref^9^ Iannone et al., 2019; ref^10^ Cryan et al., 2020). A growing body of evidence suggests that the GM may have a direct or indirect impact on central processes by activating the immune system (e.g. through inflammatory cytokines and chemokines) (ref^11^ Blander et al., 2017) and producing neurotransmitters (e.g. serotonin, gamma-aminobutyric acid, and glutamate), short-chain fatty acids (ref^12^ Aho et al., 2021), and important dietary amino acids such as tryptophan and its metabolites (ref^13^ Cryan et al., 2012). Conversely, the brain can influence gut peristalsis and sensory and secretion function primarily through the vagus nerve (ref^14^ Bonaz et al., 2018).

Publications from the Helsinki cohort have previously reported several findings regarding the relationship between GM and PD. These include a decrease in the abundance of Prevotellaceae in the GM of PD patients, a positive correlation between the relative abundance of Enterobacteriaceae and the severity of certain PD motor symptoms, a persistence of these GM changes after a 2-year period, and a connection between GM and the progression of PD (ref^2^ Scheperjans et al., 2015, ref^15^ Aho et al., 2019).

Other components of the GM comprise the mobile genetic elements (MGEs), such as plasmids and phages, which remain mostly undetermined with regard to PD. Plasmids are commonly found in bacteria and can carry a variety of genetic material, including genes related to antibiotic resistance. Phages are highly variable and are engaged in an ongoing battle with bacteria, thus affecting microbiota composition. Investigating the phage populations that may impact GM composition is important for understanding the gut microbiome and its relationship to health (ref^16^ Breitbart et al., 2003; ref^17^ Minot et al., 2011; ref^18^ Minot et al., 2012; ref^19^ Minot et al., 2013). The human virome has been linked to several illnesses, such as cancer (ref^20^ Hannigan et a. 2018), type 2 diabetes (ref^21^ Yang et al., 2021) and Inflammatory Bowel Disease (IBD) (ref^22^ Liang et al., 2021), and also to IBD-associated immunomodulation (ref^23^ Adiliaghdam et al., 2022). Interestingly, viral transfer to the gut has been tested as a treatment in a model system (ref^24^ Draper et al., 2020). Thus, research of MGEs is crucial in the context of gut microbiome and health, especially in the case of PD.

The heterogeneity of gut phages and viruses makes it difficult to determine the gut phage population. Unlike bacterial communities, which can be studied using a common set of primers targeting 16S rRNA gene sequences, there are no common primers that can be used to examine the entire phage/virus community. Nevertheless, the Gut Phage Database (GPD) (ref^25^ Camarillo-Guerrero et al., 2021) can be effectively applied to analyze gut phage populations by aligning their sequences with metagenome sequences.

Clustered Regularly Interspaced Short Palindromic Repeats (CRISPR) loci, found in bacterial genomes and plasmids, serve as a defense mechanism against MGEs in bacteria and archaea. The CRISPR system can be used to trace the history of phages encountered by bacteria in the GM. The loci consist of direct repeats (DRs) and spacers, which are complementary to phage or plasmid genetic material and allow the CRISPR-associated (Cas) system to mount a specific defense against invading MGEs (ref^26^ Hille et al., 2016). Investigation of these sequences can help to identify the phage populations encountered by bacteria. It is noteworthy that some phages carry CRISPR genes in their genomes that can target other MGEs or host genes. Even phages lacking the cas gene have been reported to have mini-CRISPR arrays (ref^27^ Seed et al., 2013; ref^28^ Medvedeva et al., 2019).

We opted to employ these methods to compare the fecal phage and plasmid populations of PD patients and control subjects, aiming to identify any differences that could potentially impact GM composition. Previous shotgun metagenome sequencing studies have already investigated the connection between PD and the gut microbiome, and our results will serve as a valuable addition to this body of research (ref^29^ Bedarf et al., 2017; ref^30^ Wallen et al., 2022; ref^31^ Qian et al., 2020).

In our analysis, we observed a diverse MGE community in the fecal samples, consistent with previous descriptions of the fecal microbiome. Notably, we found differences in alpha and beta diversity indices of plasmids between the PD and control groups. Phage communities were also highly variable, with several phages exhibiting differential abundance between the PD and control groups. Our findings add a crucial aspect to understanding the relationship between the gut microbiome and PD, highlighting the role of MGEs in this context.

## Methods

Supplementary Fig. S1 provides a summarized visualization of the methods pipeline.

### Study populations

This case-control study compared 68 patients with a diagnosis of PD according to the Queen Square Brain Bank criteria with 68 sex- and age-matched control subjects. (ref^32^ Berardelli et al., 2013) (Table 1). Exclusion criteria covered a broad range of conditions and medications that could independently affect the fecal microbiome (ref^2^ Scheperjans et al., 2015).

**Table 1.** Demographics of 136 participants in this case-control study.

| Demographics | Patients | Controls |
| --- | --- | --- |
| n | 68 | 68 |
| Female subjects | 50% | 50% |
| Age (mean ± SD) | 65.35 ± 5.42 | 64.60 ± 6.87 |
| BMI (kg/m<sup>2</sup>,<br>median [IQR]) | 27.16[24.25-29.36] | 26.21[24.09-28.05] |

### Stool sampling

Stool samples were collected at home by all PD patients and controls included in the Helsinki cohort using DNA Stool Collection Tubes containing DNA Stabilizer from the PSP® (Pre-analytical Sample Processing) Spin Stool DNA Plus Kit (STRATEC Molecular®), and were subsequently frozen and stored at −80 °C. Stool samples were shipped on dry ice to the DNA Sequencing and Genomics Laboratory at the Institute of Biotechnology, University of Helsinki, Finland for, DNA extraction, sequencing and data analysis. (ref^2^ Scheperjans et al., 2015, ref^15^ Aho et al., 2019)

### DNA extraction

DNA extraction of the samples was performed using the STRATEC Molecular® PSP Spin Stool DNA Plus Kit compatible with the DNA Stool Collection Tubes with DNA Stabilizer according to the manufacturer’s instructions. Samples were randomized between extraction batches to minimize batch effects.

### Library preparation and sequencing

After total DNA extraction, DNA libraries were prepared using the Nextera Library Preparation Biochemistry (Illumina). Shotgun sequencing was performed with the Illumina platforms Nextseq 500 (170 bp + 140 bp) and NovaSeq 6000 (150 bp + 150 bp).

### Sequencing analysis

Adapter sequences (R1: AGATCGGAAGAGCACACGTCTGAACTCCAGTCAC, R2: AGATCGGAAGAGCGTCGTGTAGGGAAAGAGTGTAGATCTCGGTGGTCGCCGTA TCATT) were first removed from paired end reads (R1 & R2) and then reads were trimmed using Cutadapt (v1.8.1) (ref^33^ Martin et al., 2011) with default parameters. Quality trimming was set at 20, accepting reads longer than 20 bp and systematically removing the first five bases. Trimmed reads were mapped against human reference sequence (GRCh38.p11) using BWA mem (v.0.7.12-r1039)(ref^34^ Li et al., 2009) and unmapped reads were assembled independently, sample by sample, using SPAdes (v3.11.1)(ref^35^ Nurk et al., 2017, ref^36^ Nurk. et al., 2013) with meta option. The final sequence data set (without the blanks) consisted of 8 029 959 916 good quality reads.

### Analysis approaches for plasmids

Reconstruction of plasmids from the metagenomic assemblies was performed using the PlasX tool (ref^37^ Yu et al., 2022). For each sample assembly, genes were called using Anvio, with “anvi-export-gene-calls --gene-caller prodigal”. Then, genes were annotated using PlasX de novo gene family database with “plasx search_de_novo_families” command. Further, COG and Pfam annotation was done using Anvio with “anvi-run-ncbi-cog” and “anvi-run-pfams” commands. COG14 and Pfam_v32 databases were used since PlasX models are compatible with only these databases. Then, “plasx predict” from PlasX was used to identify plasmid contigs from the assemblies. The threshold of 0.5 plasx score was used for the selection. The metagenomic reads were then mapped back to predicted plasmid contigs using Bowtie2 v2.4.4. If there were more than nine reads mapped with outward oriented pairs, we assumed that the contig is circular. MobMess (ref^37^ Yu et al., 2022) was used to identify plasmid systems and classify the type of the plasmid (backbone, fragment, compound). The clustering of the plasmid sequences was done using MMseqs2 v92deb92 with default settings.

We also checked the antimicrobial resistance genes within the plasmid contigs. First genes were annotated using Prokka (ref^38^ Seemann, 2014). To use the same version as our Antibiotic Resistance Genes (ARGs) study (Henttonen et al., in preparation), we used the following version of deepARG downloaded from the project git repository (git commit “ab05670”; https://bitbucket.org/gusphdproj/deeparg-ss/src/ab0567032235b4c67a36a0bc27d80fc2dd4f7eda/). Then, deepARG was run with default settings using plasmid contig genes.

To predict the COG groups of the plasmid genes, we ran reCOGnizer (ref^39^ Sequeira et al., 2022). We also predicted the taxonomy of the plasmid contigs using Kaiju (ref^40^ Menzel et al., 2016). NCBI RefSeq plasmid database (April 2022) was used to build the Kaiju database using “kaiju-makedb -s plasmids”. Plasmid contigs were thenannotated using Kaiju with options “-a mem -m 14 -s 80”.

### Phage identification using CRISPR

To identify the phage population component of the GM we decided to target CRISPR loci present in the bacterial genomes. Identification of CRISPR loci is performed with PILER-CR (ref^41^ Edgar et al., 2007). Spacers sequences are used in the CRISPR Cas system to target specific phages and protect the bacteria against them. Spacers are thus DNA sequences that allow taxonomic identification of phages. PILER-CR enables CRISPR loci identification by recognition of an area composed of highly conserved short repeat sequences (DR) interspaced with similar size sequences (spacers). We wrote bash script to obtain spacer sequences from the PILER-CR report. Each spacer sequence was then used to query Genbank-Phage 237 and RefSeq-Plasmid 200 databases using the National Center for Biotechnology Information (NCBI) Basic Local Alignment Search Tool for nucleotide (BLASTn) with default parameters. This step allowed us to assign a phage to each spacer in order to identify the different phage populations and the abundances from each biological sample.

### Phage identification using GPD

Phage identification using alignment can be challenging due to the low determination and understanding of this population and its plasticity. However, some databases regroup the known GM phage population, their classification and known hosts. This is why we decided to align the reads resulting from our shotgun sequencing against the Gut Phage Database (GPD) (ref^25^ Camarillo-Guerrero et al., 2021), using Bowtie2 v2-2.3.4.3 (ref^42^ Langmead et al., 2012) with default options.

### Statistical analysis

All statistical analyses and data visualization were performed with the R statistical programming language (v3.6.1) (ref^43^ Team RC, 2013). Quantitative variables are presented as mean and standard deviation. All p-values are double-tailed, with statistical significance accepted at an alpha of 0.05. Non-rarefied data without singletons (which could be PCR/sequencing artifacts) were used for calculating alpha diversity indices (observed richness, Shannon index, and Inverse Simpson index). These three different indices were compared statistically between groups using the Wilcoxon rank sum test. Beta diversity, based on binary Jaccard distance, was visualized with Non-Metric Multidimensional Scaling (NMDS) and compared statistically with adonis2 (an implementation of PERMANOVA) from the vegan package (ref^44^ Oksanen et al., 2012). Differential abundance analyses were done with DESeq2 (ref^45^ Love et al., 2014), which uses Generalized Linear Models assuming a negative binomial distribution, with Benjamini-Hochberg method for multiple comparison correction.

#### Identification of viral sequences

To identify viral contigs from metagenomic assemblies, we used VIBRANT v1.2.1 (ref^46^ Kieft et al., 2020) with default settings. The taxonomic annotation of the identified contigs was done using Kaiju v1.8.2 with “-a mem -m 14 -s 80” settings and Kaiju “viruses” database (downloaded on March 2022). Identification of CRISPR-Cas systems in the viral contigs was performed using CRISPRCasTyper v1.8.0 (ref^47^ Russel et al., 2020) with options “--minNR 2 --prodigal meta”. We used SpacePHARER v5-c2e680a (ref^48^ Zhang et al., 2021) to identify targets of the spacers.

#### Identification of crAss-like phages

To specifically identify crAss-like phages from our viral contigs, we performed the same methods as described in Guerin et al., (2018) (ref^49^). The proteome of the viral contigs was aligned against conserved crAss-like phage proteins; UGP_018 and UGP_092 (crAss polymerase, and crAss terminase, respectively). The alignment was done using BLAST v2.12.0 (ref^50^ Camacho et al., 2009) with e-value threshold 1E-05. If there was a blast hit with query alignment length ≥350bp, we selected them as putative crAss-like phages. We further filtered the results by selecting only contigs larger than 70kbp.

### Genetic code diversity and coding density calculation

We predicted genes of the phages using prodigal with three different runs using “-g 4”, “-g 11”, and “-g 15” to get gene prediction with genetic codes of 4, 11, and 15 respectively. The predicted genes were then used to calculate coding density. The coding density was calculated with the same method as Borges et al., (2022)(ref^51^). using “get_CD.py” python code from https://github.com/borgesadair1/AC_phage_analysis (downloaded August 2022).

#### Machine learning model

We studied whether phages of interest could be used to predict the classification of samples into PD or the control group utilizing a Random Forest machine learning approach (ref^52^ Pedregosa et al., 2011). Using the relative abundance of 60 phages resulting from the selection of 20 phages from the three following groups, we ran three separate analyses for different phage groups: lowest adjusted p-value, highest log2 fold change with significant adjusted p-value, and highest base mean with significant adjusted p-value, with base mean representing the average of the normalized counts of all samples dividing by size factors. The analyses were used to select the four phages with the highest predictive potential. By choosing only the top four phages from each analysis we aimed to build a final model that uses a limited number of potential biomarkers instead of the whole GPD dataset. For each run, we divided our cohort into two groups: a training group (75% of the total dataset) and a test group (the remaining 25%) randomly selected, but evenly distributed among PD and control groups. Random Forest model (non-parametric supervised learning method) was created using 200 trees (n_estimators=200) and entropy criterion parameters (parameters selected using GridSearchCV). To introduce bootstrapping to the dataset splitting step, we split randomly the original dataset 1000 times into a train and test group and fit the model for each split followed by evaluation of the F1 and area under the curve (AUC) scores. The impact of variables (phage abundance) was calculated using SHAP python package with shapTreeExplainer function (ref^53^ Lundberg et al., 2020).

## Results

### Plasmids

A total of 44,409 plasmid contigs were identified from all 136 samples using the PlasX tool. The number of plasmid contigs in control samples was 24,559, while in PD samples it was 19,850, with an average of 361 plasmid contigs per control (minimum 139 and maximum 649 plasmids) and 292 plasmid contigs per PD sample (minimum 155 and maximum 662 plasmids). The average length of plasmid contigs was 4,065 bp, with a range of 1000 bp to 275,216 bp. Using MMseqs2 clustering, we clustered the 44,409 plasmid sequences based on their sequences and obtained 29,096 clusters (Supplementary Table S1, Supplementary Table S2). We also used the MobMess tool to group plasmids into plasmid systems, which are groups of plasmids consisting of backbone plasmids and compound plasmids. Backbone plasmids only contain core genes that are conserved within the group, while compound plasmids contain both backbone genes and additional cargo genes. For instance, the predicted plasmid system “ps30” includes one backbone plasmid with one backbone gene and nine compound plasmids with seven additional cargo genes (Supplementary Table S3).

A total of 561 plasmid systems (Supplementary Table S3) were identified from the 44,409 plasmid contigs. Among them, 22 592 were identified as “maximal_not_in_system”, indicating intact plasmids that were not members of any plasmid systems. Additionally, 18 749 were identified as “fragment”, 760 as “compound plasmid”, and 2,308 as “backbone plasmid” (Supplementary Table S4). To determine taxonomic annotation, the NCBI RefSeq plasmid database (April 2022) was used. *Enterococcus* and *Vescimonas* plasmids were the most commonly observed on the genus level (Supplementary Table S5, Supplementary Fig. S2).

Plasmids are a type of mobile genetic element commonly found in bacterial cells. They vary in length, copy number, and coding capacity for proteins. We compared the plasmid populations between PD and control groups, as well as factors such as gender, BMI, and age. We found significant alpha diversity differences between PD patients and control subjects using pairwise Wilcoxon rank sum test (Inv: p = 0.00080; Sha: p = 0.00076; Obs: p = 0.00071) and Spearman’s rank correlation test (Inv: p = 0.00066; Sha: p = 0.00061; Obs: p = 0.00057) (Figure 1). Additionally, we found a statistically significant correlation between BMI and plasmid alpha diversity using Spearman’s rank correlation test (Inv: p = 0.00463; Sha: p = 0.00479; Obs: p = 0.00472). No other variables showed significant differences or associations. Plasmids also displayed significant beta diversity differences between the PD and control groups (p = 0.00050) (Supplementary Fig. S3).

**Figure 1.**
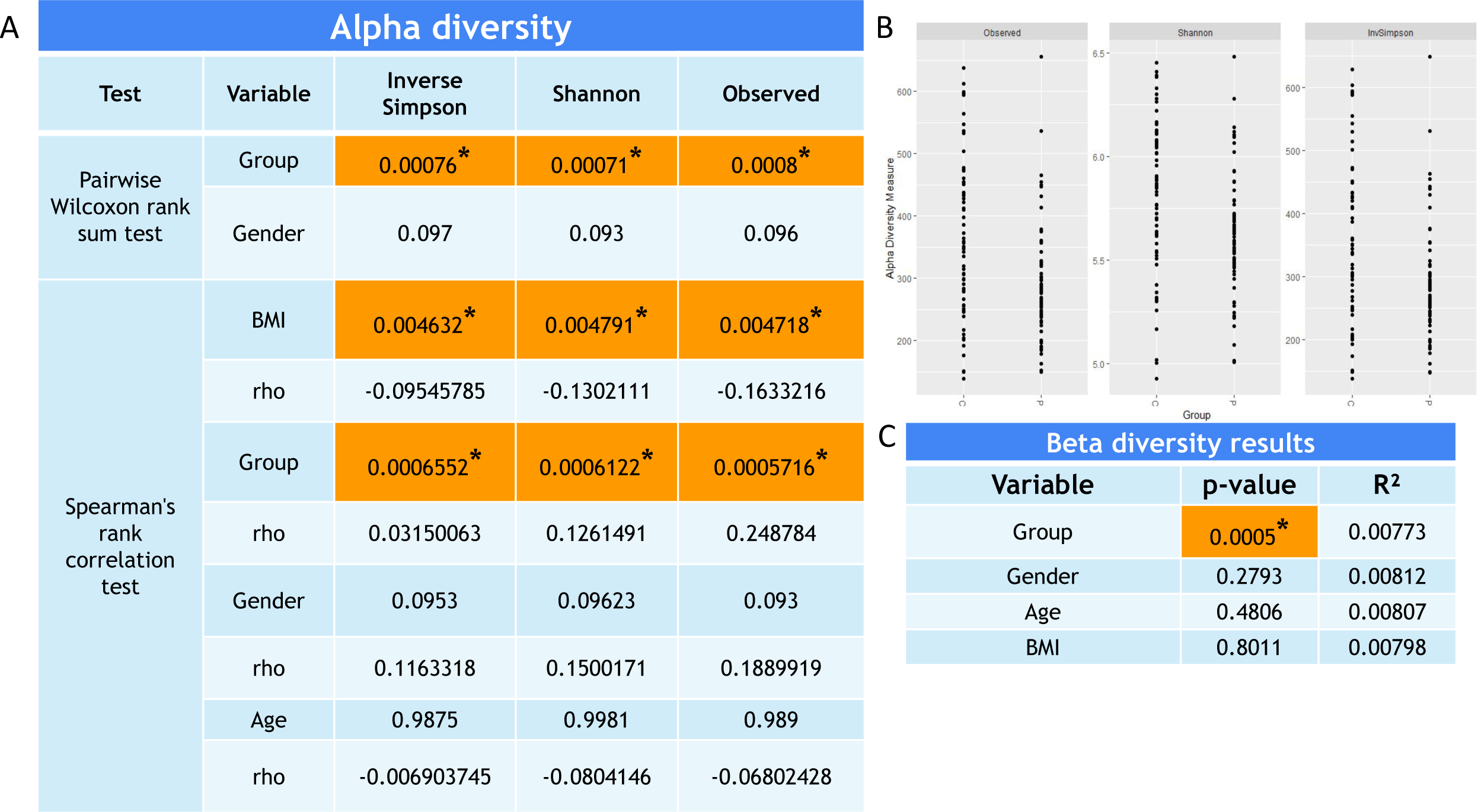
Statistical analysis result obtained for Plasmid population investigation. **[A]** Alpha diversity results using Pairwise Wilcoxon rank sum test and Spearman’s rank correlation test. **[B]** Alpha diversity measure plot for each indices for control (C) and Parkinson (P) groups. **[C]** Beta diversity results obtained using Aitchison distance in univariable PERMANOVA models.

In addition to diversity, we also checked the total abundance of the plasmids. The total abundance of plasmids did not show any significant difference between PD and control groups (Supplementary Fig. S4).

To gain general insight into plasmid gene functions, Clusters of Orthologous Genes (COG) categories and COG functions of plasmid genes were annotated. The most common COG categories were “Replication, recombination and repair”, “Cell cycle control, cell division, and chromosome partitioning”, and “Transcription”. The most common COG functions were “ParA-like ATPase”, “Site-specific DNA recombinase SpoIVCA/DNA invertase PinE”, and “Chromosome segregation ATPase Smc” (Supplementary Fig. S5, Supplementary Table S6).

We examined the gene populations carried by plasmids and found differences in alpha diversity between PD and control groups using both Wilcoxon rank sum test (Obs: p = 0.042) and Spearman’s rank correlation (Obs: p = 0.041) (Figure 2). The beta diversity also showed a difference between the PD and control groups (p = 0.027) (Supplementary Fig. S3). However, we did not find distinct activities that differed between any of our comparisons in the functional characterization of the genes.

**Figure 2.**
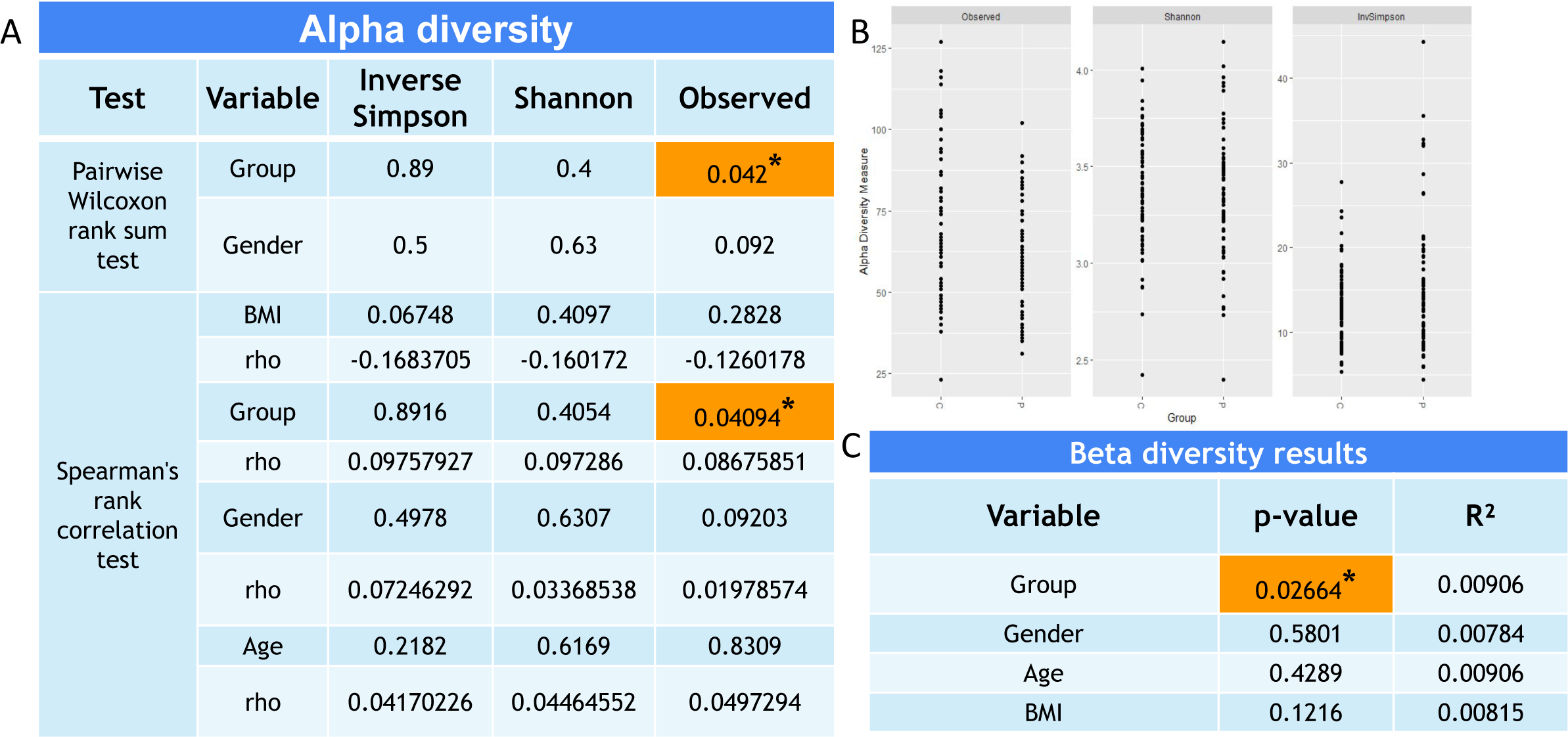
Statistical analysis result obtained for Plasmid genes population investigation. **[A]** Alpha diversity results using Pairwise Wilcoxon rank sum test and Spearman’s rank correlation test. **[B]** Alpha diversity measure plot for each indices for control (C) and Parkinson (P) groups. **[C]** Beta diversity results obtained using Aitichon distance in univariable PERMANOVA models.

We also identified antibiotic resistance genes (ARGs) within the plasmid genes. In every sample, at least one ARG was found within the plasmid genes, with vancomycin resistance gene, *vanS*, being the most frequently observed gene in this category (Supplementary Fig. S6, Supplementary Table S7). Additionally, the predominant class of antibiotic resistance genes present on plasmids in the gut microbiota of both PD and control groups was the glycopeptide antibiotic class (Supplementary Fig. S6, Supplementary Table S7).

Plasmids are the most common vehicles moving ARGs between bacterial cells and antibiotic usage might be related to PD (ref^49^ Mertsalmi et al., 2020). No differences were observed in this study between PD and controls, sex, BMI, or age for alpha diversity or for beta diversity regarding ARGs (Supplementary Fig. S7) or their classes (Supplementary Fig. S8) when evaluating plasmid contigs.

### Phages

#### De novo identification of viral contigs from metagenomic assembly

Using Vibrant, we were able to detect 111 099 viral contigs from the entire dataset. Among these, 61 793 were from c-ontrol samples and 49 306 from PD samples (Supplementary Table S8). Although most of the contigs were low quality draft, some complete circular quality contigs were also observed (Supplementary Fig. S9). The Vibrant classification revealed that the number of “lytic” type viral contigs was higher than “lysogenic” type in the gut microbiota (Supplementary Fig. S9). Additionally, we found that “lysogenic” type contigs tended to be longer than “lytic” type contigs. At the order level, Caudovirales was the most dominant order (Supplementary Fig. S10; Supplementary Table S9). At the family level, the most prevalent taxa were Siphoviridae, Myoviridae, and Podoviridae, respectively.

#### Identification of CRISPR-Cas systems in viral contigs

In our study, we focused on examining CRISPR-Cas systems within the identified viral contigs. Our analysis revealed that out of the 111,099 viral contigs examined, 144 contained cas genes; type II-D was the most frequently observed cas gene type. Overall, 163 cas genes were detected across the 144 viral contigs, with the majority (n=136, 84%) belonging to the II-D type. Other Cas gene types that were detected in the identified viral contigs included IV-B, V-F1, VI-D, I-C, VI-A, V-A, VI-B1, I-E, II-B, and I-B (Supplementary Table S10). Of the 144 viral contigs in which a cas gene was detected, 47 (33%) were identified within a metagenome-assembled genome that was constructed in our previous study (ref^54^ Duru et al., 2023). Notably, our observations revealed that among the cas genes we identified within viral contigs, only 20 had proximate CRISPR arrays, while for the majority (n=124), no nearby CRISPR arrays were detected.

Our analysis unveiled the presence of CRISPR arrays lacking cas genes within viral contigs. A total of 1621 CRISPR arrays were detected within the viral contigs, distributed across 1529 distinct viral contigs. Only 20 of them were cas gene. It is noteworthy that a majority of these CRISPR arrays consist of less than three repeats (Supplementary Table S10). We also checked whether the spacers target any known phages, and observed that targets of the spacers were mainly *Faecalibacterium* and *Escherichia* phages (Supplementary Table S10).

#### CrAss-like phages amd prophages

A more detailed search was conducted for crAss-like phages, which are the most prevalent phages found in the human gut metagenome (ref^55^ Dutilh et al., 2014) of the identified viral contigs. A total of 118 crAss-like phages were identified, with 92 containing both crAss polymerase and crAss terminase, 11 containing only crAss polymerase, and 15 containing only crAss terminase. Using presence/absence clustering, we identified four main groups for CrAss-like phages (Figure 3). However, no significant differences were observed between PD and control groups.

**Figure 3.**
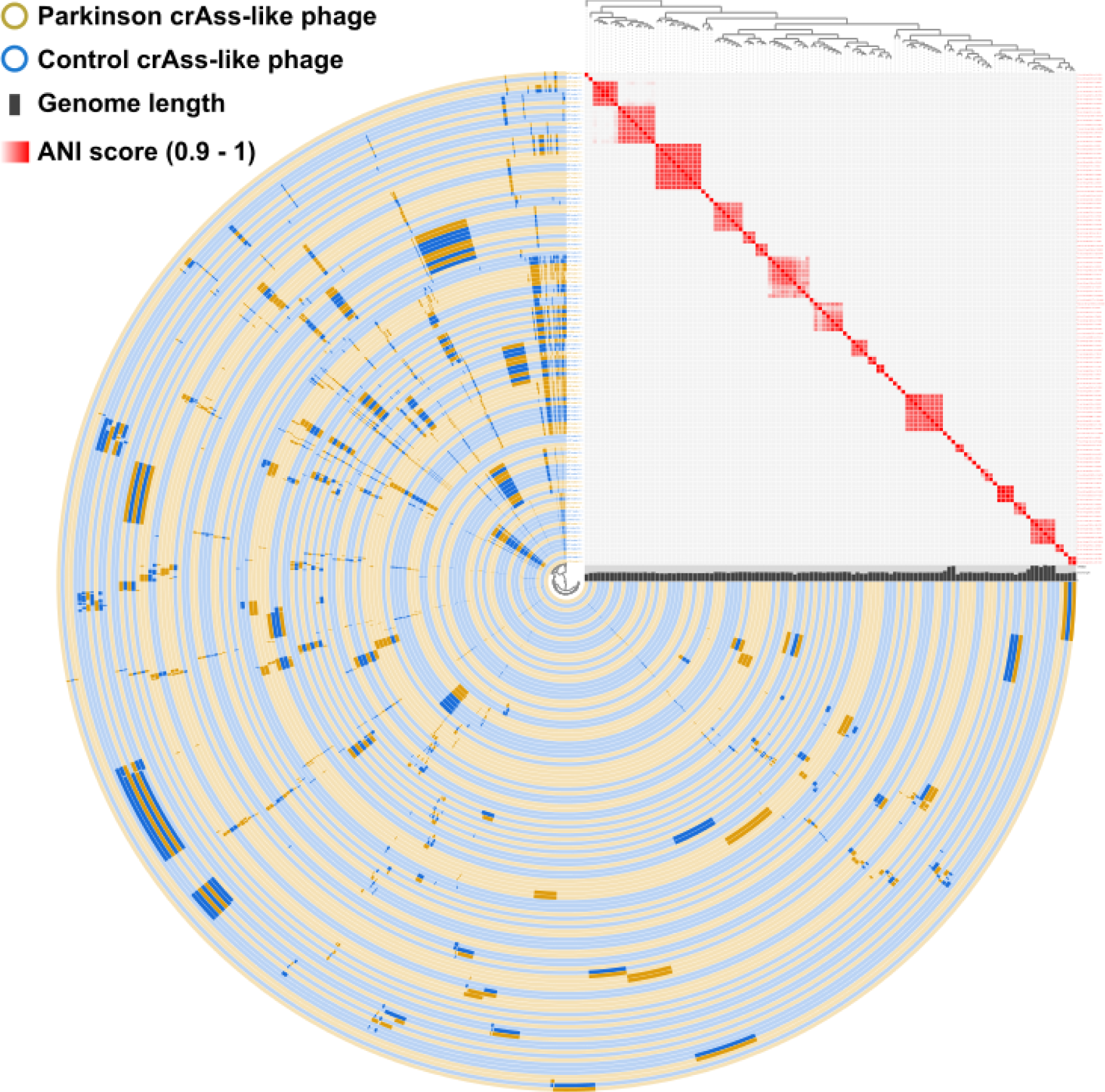
The 119 inner circular layers correspond to the 119 crAss-like phage. Phage genomes that were created from the Control group and Parkinson group are shown in blue and yellow, respectively. Phages are ordered according to gene presence/absence. Tree at the top also clusters genomes based on gene presence/absence. ANI scores are shown at the top-right corner with red gradient. The length of the MAGs are shown with grey bars.

In addition we investigated the prophages proportion in our samples. Out of our 136 samples, 6849 phages have been identified with a large prevalence of dormant prophages. Regarding the 138 actives prophages, there were no statistically significant difference between PD and control groups.

#### Alpha and beta diversity of phages using GPD

The alpha diversity of phages was lower in the PD population than in controls (Pairwise Wilcoxon rank sum test: p = 0.0021; Spearman’s rank correlation test: p = 0.0019) (Figure 4). We identified a lower observed richness (Pairwise Wilcoxon rank sum test: p = 0.0320; Spearman’s rank correlation test: p = 0.0294) in women (Figure 4). No significant associations of alpha diversity were found for age and BMI.

**Figure 4.**
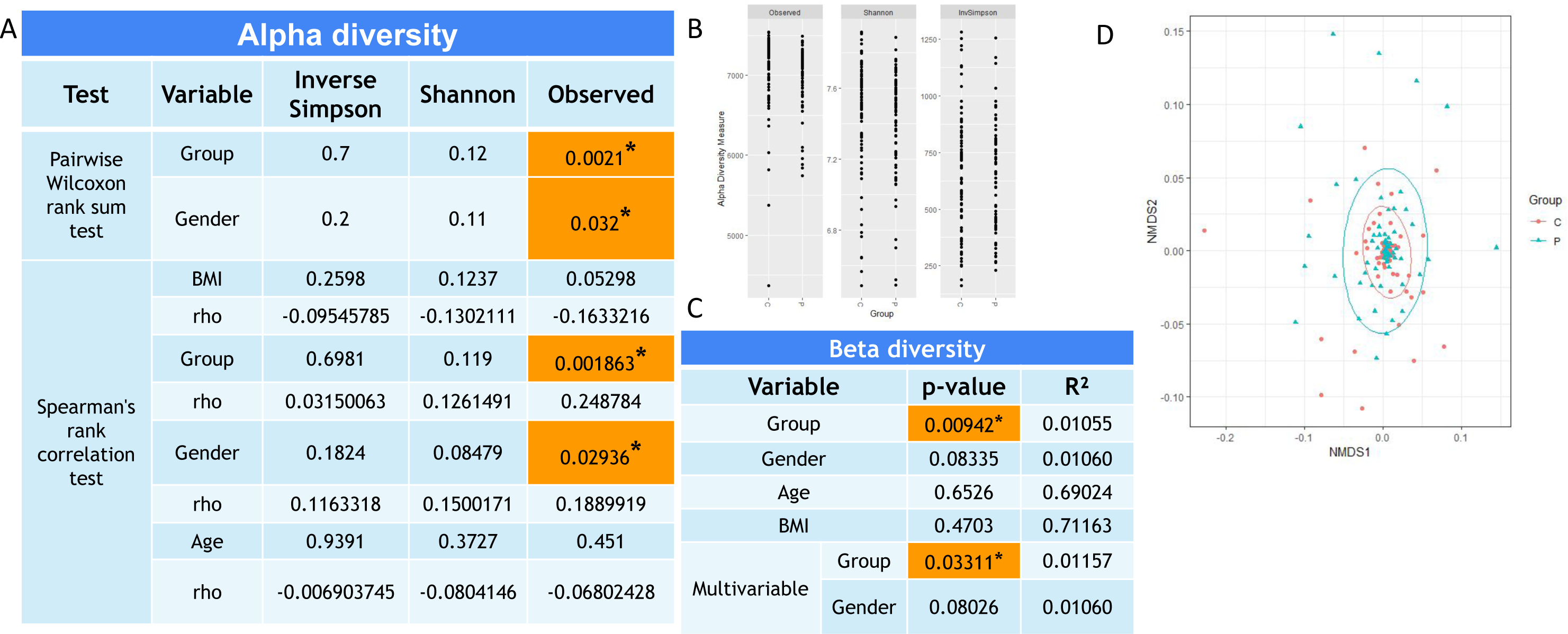
Statistical analysis result obtained for phage population investigation using GPD approach. **[A]** Alpha diversity results using Pairwise Wilcoxon rank sum test and Spearman’s rank correlation test. **[B]** Alpha diversity measure plot for each indices for control (C) and Parkinson (P) groups. **[C]** Beta diversity results obtained using Jaccard distance in univariable and multivariable PERMANOVA models. **[D]** Nonmetric Multi-Dimensional Scaling (NMDS) ordination plot of the control (C) and PD patients (P) samples. Ordination based on Jaccard distance, using non rarefied data. Ellipses indicate 95% confidence intervals. Each point represents one sample. The closer the points are to one another the more similar the phage compositions of the samples are.

Beta diversity quantifies the community dissimilarity between samples, and showed differences between patients and controls using binary Jaccard distance in the univariable analysis (i.e. single explanatory variable) PERMANOVA model (*p* = 0.00942, r² = 0.01055) (Figure 4). A low *p*-value difference (of potential interest but not statistically significant) was observed between the sexes (*p* = 0.08335, r² = 0.01060). Therefore, we decided to investigate these two variables in a multivariable model (i.e. a model with more than one explanatory variable), which confirmed a difference between patients and controls (*p* = 0.03311, r² = 0.01157), but not between the sexes (*p* = 0.08026, r² = 0.01060). Furthermore, no association was identified with age or BMI.

#### Differential abundance analysis using GPD

We utilized the DESeq2 tool to examine the difference in phage abundance between PD patients and controls, resulting in a total of 1,109 differentially abundant phages (Supplementary Table S11). To narrow our focus, we selected a subset of statistically significant phages, including 20 from three groups: those with the lowest adjusted p-value (Supplementary Table S12), the highest log2 fold change (Table 2), and the highest base mean (Supplementary Table S13). For the highest log2 fold change group, we divided it into two subgroups to identify phages with the largest positive and negative log2 fold changes.

**Table 2.**
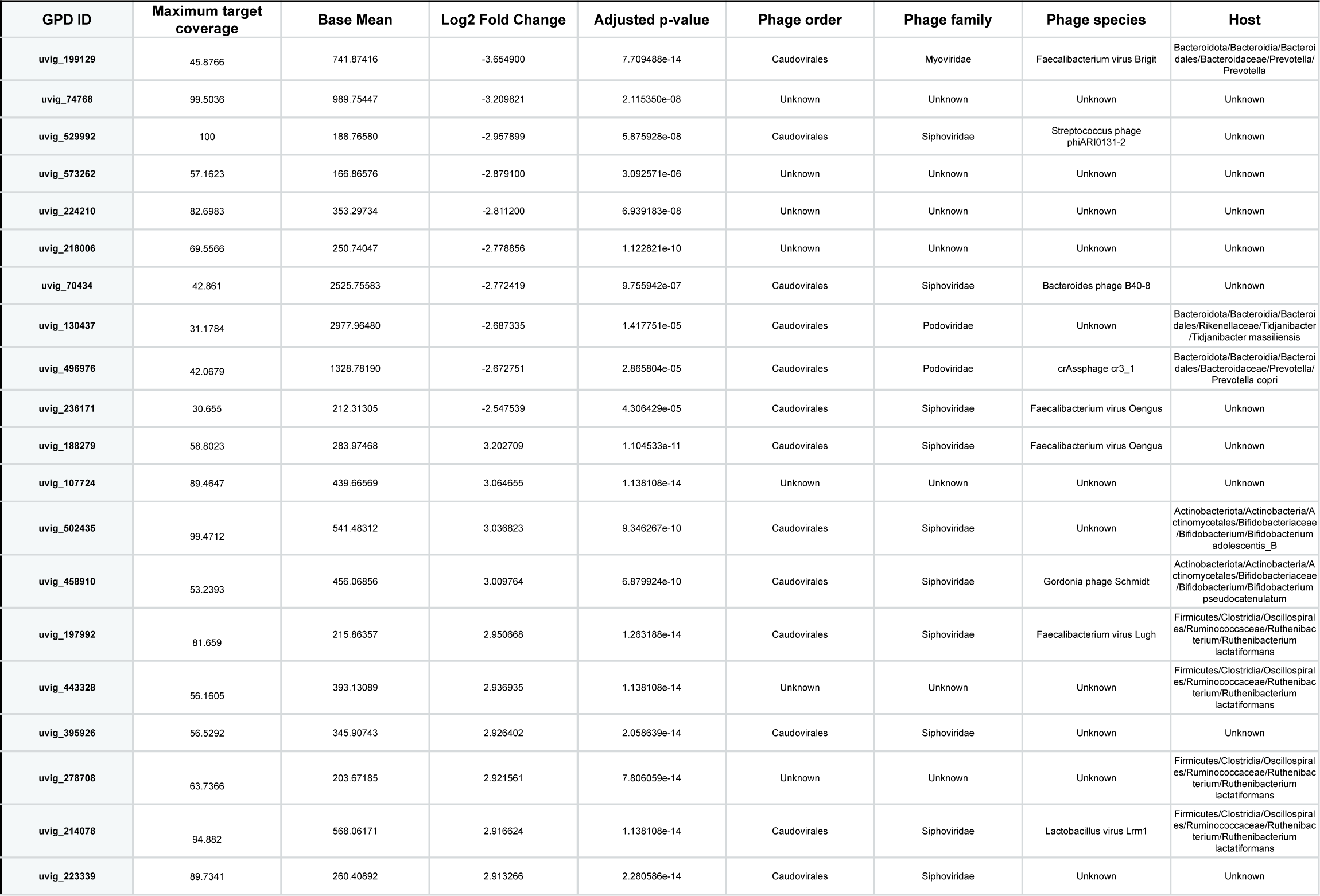
20 significantly differentially abundant phages with the highest Fold change obtained using the GPD approach. The first 10 phages present a negative Fold change and the 10 last phages present a positive Fold change.

In terms of phage classification, the majority of the selected phages (42/60) belong to the Caudovirales order, with most of them falling under the Siphoviridae (27 phages) or Myoviridae (9 phages) family. Only four phages were classified into other families (three Podoviridae, one Herelleviridae), and one of the selected Podoviridae phages is also a representative of the common crAssphage. Hosts were identified for 27 phages, with most of them having similar hosts among several bacterial genera, including *Ruthenibacterium* (13 phages), *Bifidobacterium* (6 phages), and *Prevotella* (3 phages). Other identified hosts were *Tidjanibacter* (2 phages), *Faecalibacterium* (1 phage), *Roseburia* (1 phage), and *Lachnospira* (1 phage). To gain a better understanding of gene functions in selected phages, we annotated the gene functions in the phages selected based on log2 fold change (Figure 5). We found phage-related genes coding for tail and capsid protein, as well as other genes of interest such as those coding for virulence protein and metallo-beta-lactamase.

**Figure 5.**
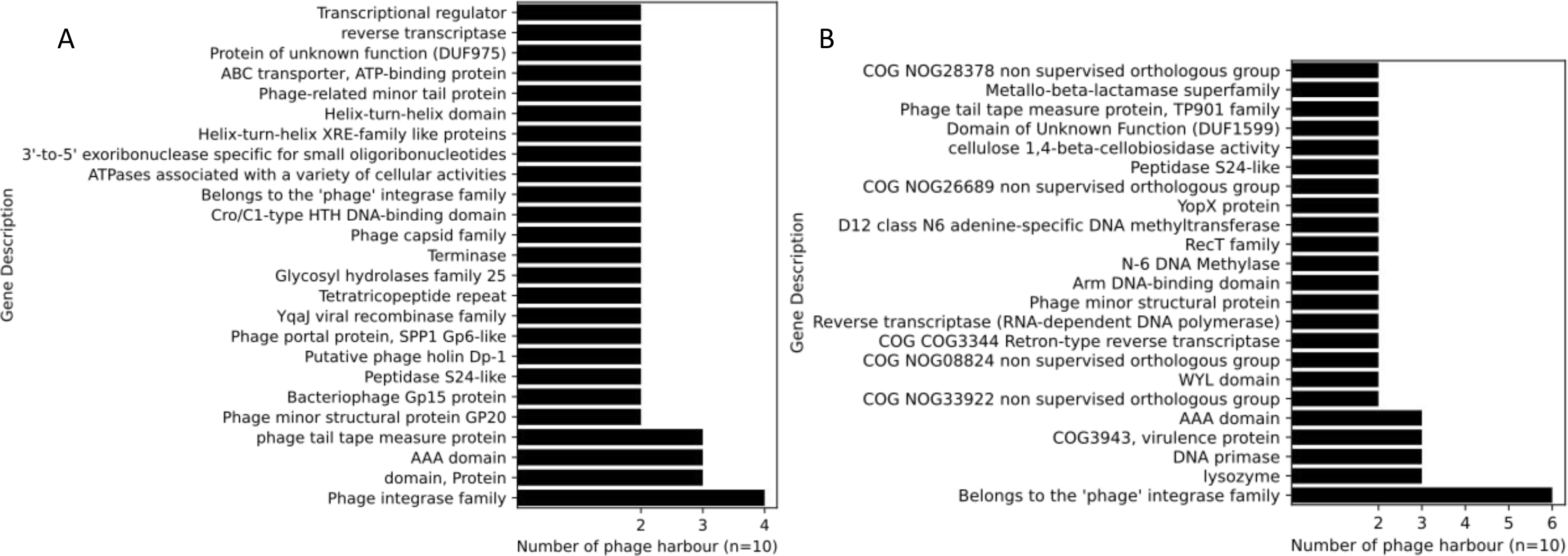
Gene composition of phages significantly differentially abundant and presenting the largest fold change between PD and Control. [A] Gene composition of phages with a positive log2Fold change. [B] Gene composition of phages with a negative log2Fold change.

In addition to the species specific abundance analysis, we also compared total phage abundance between PD and control groups. The analysis did not show any significant difference between groups (Supplementary Fig. S4).

#### Alpha and beta diversity using CRISPR data

The phages identified using CRISPR did not show significant dissimilarity between PD and control groups regarding alpha and beta diversity and differential abundance. However, alpha diversity was significantly associated with BMI as assessed with Inverse Simpson (Supplementary Fig. 11).

#### Machine learning model

A machine learning approach was used to investigate whether the relative abundance of selected phages could be used to predict sample classification into either PD or control groups. The final model for Random Forest was created using the selected phages from the three groups (lowest adjusted p-value, highest log2 fold change, and highest base mean), resulting in 11 phages (one phage was redundant in two groups) (Figure 6 B).

**Figure 6.**
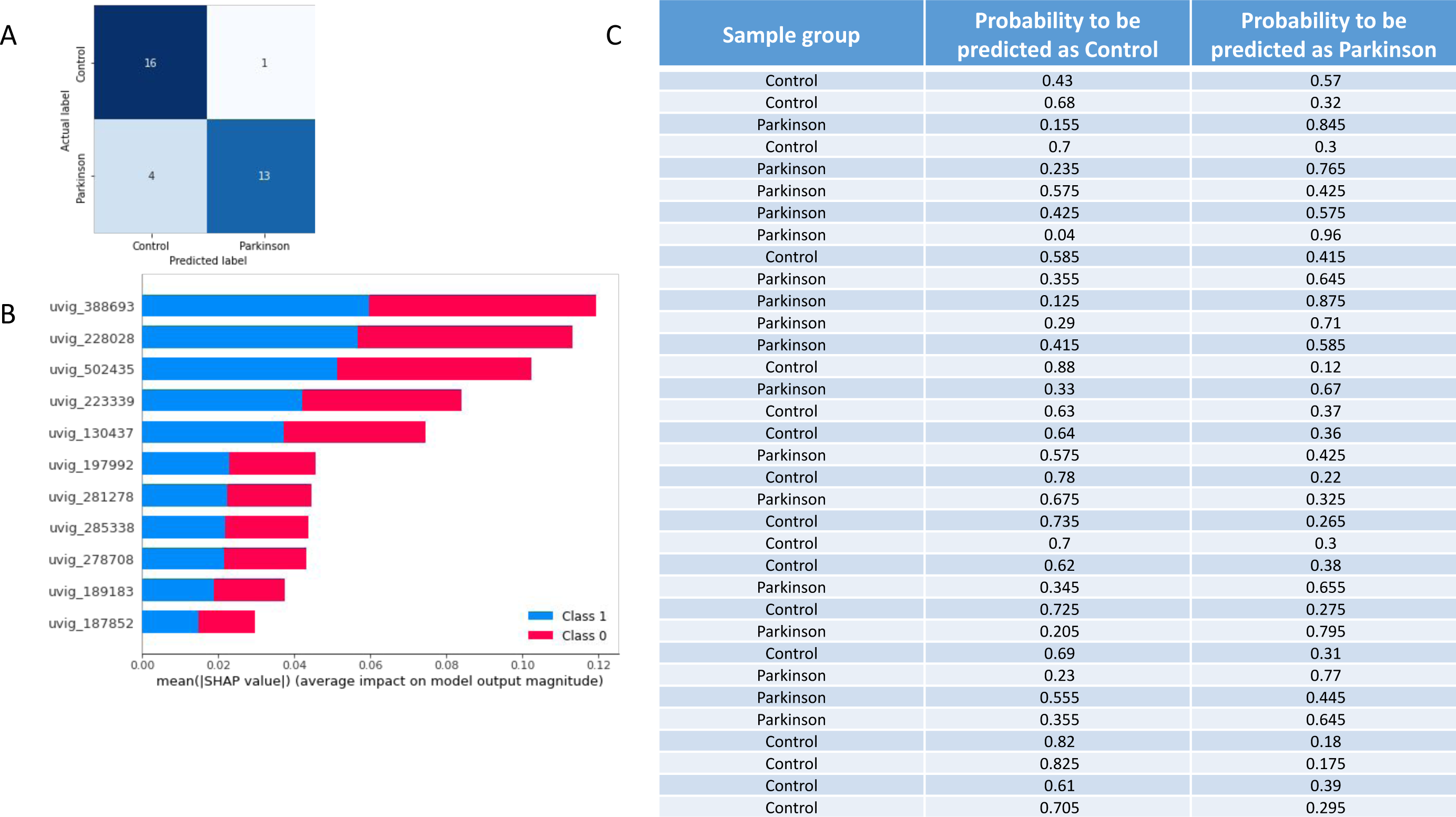
[A] Confusion matrix presenting the classification performance of our machine learning model. [B] Feature importance plot generated by SHAP TreeExplainer. The plot provides the impact of each features on the model predictions. The stacked bars indicate the mean magnitude of the SHAP values for each feature (phage relative abundance) across all samples in the dataset. (class 1:Parkinson and class 0:Control).

The 1000 models generated an average AUC with a 95% confidence interval of 0.737-0.746. The top-performing model achieved an AUC score of 0.969 and an F1 score of 0.839. Using this model, we were able to accurately predict 16 of 17 samples from the control group and 13 of 17 from the PD group in the test dataset, (sensitivity=0.76, specificity=0.94) (Figure 6 A). Furthermore, the model’s prediction probability allowed for easy identification of incorrectly classified samples (Figure 6 C).

#### Genetic code diversity in phages

Recently, it has been shown that phages can adapt to use alternative genetic codes (other than standard code (code 11)) to increase their adaptability (ref^51^ Borges et al., 2022) by recoding the stop codon to prevent premature production of late stage proteins. We tested the coding density of phages using standard code, code 4, and code 15, which are the three codes known to provide a recoded stop-codon. Of the 142 809 GPD phages, 139 316 (97.5%) phage genomes were predicted to be using standard code, 2,872 (2.01%) using code 15, and 621 (0.43%) to be using code 4 (Supplementary Table 14). For the phages using code 4, the average coding density was 90%, while for the phages using standard code the corresponding figure was 70%. For phages using code 15, the average coding density was 91%, while for phages using standard code it was 63%. This indicates that, for some phages, structural gene annotation can be significantly improved by using a different genetic code during the annotation run.

We also checked the coding density of phage contigs that were identified using Vibrant. Of all 111 099 phage contigs, 107 270 (96.5%) were predicted to be using standard code, 996 (0.89%) code 15, and 2,830 (2.54%) code 4 (Supplementary Table 14). Unlike for GPD phages, code 4 was the second most common genetic code preference in our phage contigs.

## Discussion

Viruses and phages constitute the most diverse and extensive group of biological entities, along with other MGEs like plasmids. They provide an adaptable mechanism for horizontal gene transfer, enabling the movement of genetic material even between distantly related organisms. Viruses and phages have a significant impact on bacterial and archaeal communities, either by killing them or by enhancing their fitness through the introduction of new genes into their genomes. Previous research has established an association between Parkinson’s disease (PD) and gut microbiota (GM) composition at the family, genus and species level, which has been linked to specific metabolite production (ref^2^ Scheperjans et al., 2015, ref^56^ Parashar et al., 2017, ref^57^ Hopfner et al., 2017, ref^30^ Wallen et al., 2022). Therefore, it is important to investigate the nature of these associations and possible causes for GM alterations in PD patients. MGEs have the potential to alter the gut microbiota, and studying them could help us understand the community dynamics and develop new approaches to control the gut environment. As a result, the focus of gut microbiota research has started to shift towards MGEs.

A previous study reported no significant difference in the abundance of prophages and plasmids between PD and control samples, but the total virus abundance was decreased in the PD group (ref^29^ Bedarf et al., 2017). However, another study noted a significant difference in prophage and plasmid populations between PD and control groups based on Shannon and Chao1 indices, and total prophage abundance was found to be higher in the PD group than in the control group (ref^58^ Mao et al., 2021). Therefore, further investigations are required to understand the relationship between the MGE population in the gut and PD. In this study, we examined the differences in MGE populations between PD patients and control subjects.

We investigated two types of MGEs, namely plasmids and phages, which have been shown to play a crucial role in regulating the bacterial population of GM and facilitating horizontal gene transfer among bacteria (ref^59^ Suzuki et al., 2019, ^60^ Broaders et al., 2013). The occurrence of horizontal gene transfer between phylogenetically distant bacterial groups highlights the importance of the ecological environment as a driving factor for gene exchange (ref^61^ Kurokawa et al., 2007; ref^62^ Smillie et al., 2011). Our findings revealed notable differences in the community of these mobile elements in PD and control groups. Notably, the variation in GM composition is known to be influenced by various factors, such as BMI (ref^63^ Liu et al., 2021; ^64^ Odamaki et al., 2016), which can impact MGEs. To account for possible confounders, including sex, age, and BMI, we adjusted our statistical models, but residual confounding cannot be entirely excluded.

Plasmid populations and plasmid encoding elements reflect the association and co-evolution of hosts and bacteria. Plasmid populations have been investigated at both diversity and gene levels. In this study, we were interested in the plasmid composition of the GM and the genes carried by plasmids. Our analysis revealed differences in alpha and beta diversity of plasmids between PD patients and controls, which is in agreement with an earlier study (ref^58^ Mao et al., 2021) that found significant beta diversity differences for plasmids between these two groups. However, there was no significant difference in the total abundance of plasmids observed between the PD and control groups, consistent with the previous findings (ref^58^ Mao et al., 2021).

Phages are highly variable and can differ substantially between individuals, similar to bacteria. We examined phage communities in PD and control groups using two different approaches for alpha and beta diversity analyses. The first approach involved investigating CRISPR loci, which can provide insights into the phage populations encountered by the gut bacterial community and the ecological competition between phages and bacteria. The second approach involved aligning reads to the Gut Phage Database (ref^20^ Camarillo-Guerrero et al., 2021). Only the second approach revealed statistically significant differences between groups, indicating that alpha and beta diversity of the phage population differed between PD and control subjects.

Furthermore, previous research has shown differences in phage diversity between individuals with PD and healthy controls, providing further support for our observed dissimilarities in alpha and beta diversity between the two groups. Plasmids carry genes involved in host-microbe and microbe-microbe interactions, such as virulence genes and ARGs, which can impact the GM population (ref^56^ Broaders et al., 2013). Our results indicate that the plasmid gene composition differs between PD patients and controls, as demonstrated by the rank sum test (p = 0.042), Spearman’s rank correlation (p = 0.041), and beta diversity (p = 0.027). In addition to general functional annotations, we identified ARGs within the plasmid genes, with the *vanS* gene being the most prevalent. This gene plays a crucial role in vancomycin-induced resistance, a member of the glycopeptide class of antibiotics (ref^61^ Marshall et al., 1998). Vancomycin has been shown to substantially decrease gut microbiota diversity (ref^62^ Vrieze et al., 2017). The high frequency of the *vanS* gene on plasmids in the gut environment may reflect an adaptation process by gut bacteria to vancomycin resistance. However, our analysis of plasmid-borne ARGs did not reveal any significant difference in diversity between PD and control groups.

We aimed to identify possible phage genomes that would be differentially abundant among the PD group and the control group. Therefore, in addition to the diversity analysis, we also performed differential abundance analysis using the GPD database. Our results indicated that 1,109 phages were differentially abundant between PD and controls. It is noteworthy that a previous investigation reported no significant difference in prophage abundance between these two groups (ref^29^ Bedarf et al., 2017). However, we recognized that database selection is a critical factor in assessing phage abundance. Specifically, we employed the Gut Phage Database (GPD) (ref^25^ Camarillo-Guerrero et al., 2021), which encompasses 142 809 non-redundant gut phages. In contrast, the earlier study (ref^29^ Bedarf et al., 2017) relied on a database of 760 phages. This marked disparity in database size and scope likely accounts for the differences observed between our study and the previous investigation. Another study, in turn, reported 241 significantly differentially abundant virus OTUs between two groups using a database with 45 033 dereplicated virus OTUs (ref^65^ Tisza et al., 2021).

Based on our differential abundance analysis, the phages with the most interesting differences between PD and control subjects belong to the other Caudovirales. Phages belonging to this order are expected to be found in the human gut and represent the most abundant phages in the GM (ref^65^ Tisza et al., 2021). Phage identification using homology is a challenging approach due to the lack of taxonomical information in available databases. The database used in this study is mainly composed of machine learning predicted phages for which classification and host prediction are not systematic (ref^25^ Camarillo-Guerrero et al., 2021). Moreover, phages are evolving in parallel with their hosts; it is therefore difficult to have a complete and up to date database containing taxonomically classified phages and their associated hosts. The phages identified in our study are only double strand DNA phages and prophages. Investigation of single strand DNA phages and RNA phages would require different sampling approaches and are not included in this study. However, 95% of GM phages are known to be non-enveloped tailed double strand DNA phages or Caudovirales, which is consistent with our results (ref^66^ Sausset et al., 2020). Our study represents a proxy of the gut microbiota due to the sampling method used, but such approximation is necessary to avoid invasive sampling methods such as surgery or biopsy.

Due to the large number of differentially abundant phages that we found (n=1 109), we decided to narrow our focus to a subset of 60 phages. We selected the top 20 phages based on the lowest adjusted p-value, highest log2 fold change, and highest base mean from the three groups. These phages belonged to several taxonomic groups, including Caudovirales, Siphoviridae, Myoviridae, Podoviridae, and Herelleviridae, consistent with a previous study (ref^63^ Tisza et al., 2021). Our analysis of phage host information indicated that the selected phages commonly target bacterial genera such as *Bacteroides*, *Faecalibacterium*, and *Prevotella*, which are prevalent in the gut microbiome and have been linked to PD pathology in earlier studies (ref^65^ Shkoporov et al., 2019, ref^66^ Shkoporov et al., 2018). These findings provide further support for the potential use of these phages to manipulate the gut microbiome and alleviate PD symptoms.

We utilized the relative abundance data of 60 selected phages to examine the feasibility of constructing a predictive model for the detection of PD based on phage abundance data. The model performance was evaluated by calculating its area under the curve (AUC) to be approximately 0.75. Consequently, we decided to reduce the number of phages used in the model creation process in order to identify potential biomarkers with greater specificity. Finally, we generated 1000 models utilizing only the abundance of 11 phages. The resulting models yielded a 95% confidence interval for the average AUC between 0.737 and 0.746. The optimal AUC score obtained was 0.969. Interestingly, the virome of gut microbiota has been used in an earlier study (ref^65^ Tisza et al., 2021) to create a predictive model for the detection of PD, and the model was reported to attain an AUC score of 1.00 (ref^65^ Tisza et al., 2021). It should be noted that before this study, the same samples employed here were used to create a predictive model using 16S bacterial community data, which yielded a slightly better score than the model created here (ref^2^ Scheperjans et al. 2015).

We computed the overall phage abundance of samples and then compared the PD and control groups based on comparison to the GPD database. Our findings reveal that no significant disparity exists between the two groups. Nevertheless, prior investigations have presented diverse outcomes regarding the overall phage abundance. Specifically, Bedarf et al. (2017)(ref^29^) reported a decrease in total virus abundance in the PD group, while Mao et al. (2021)(ref^58^) observed an increase in total virus abundance in PD. Hence, we conclude that phage specific abundance analysis may be more informative than the total phage abundance.

We also conducted a prediction of viral contigs within the metagenomic assemblies, which led to the discovery of 111 099 viral contigs. Upon identification, it was observed that “lytic” type viral contigs were more common than “lysogenic” type viral contigs in the gut microbiota of our samples. Previously a gut virome study also reported that “lysogenic” type viruses do not dominate the gut ecosystem (ref^67^ Bhardwaj et al., 2022). In our earlier study, we found that about 28% of the predicted viral sequences are actually part of microbial genomes (ref^54^ Duru et al., 2023). Possibly, the high number of “lytic” viruses might explain why there were relatively few phages within the microbial genomes. Taxonomic annotation of the predicted viral contigs indicated that Caudovirales is the dominant order, which is in line with the results of previous gut virome studies (ref^25^ Camarillo-Guerrero et al., 2021, ref^65^ Tisza et al., 2021). Our viral contig sequences could be useful for constructing future databases. Additionally, we have detected CRISPR-Cas systems within the viral contigs. Specifically, of the 111 099 viral contigs, only 144 contained cas genes, and of these 144, only 20 possessed both CRISPR array and cas genes. In addition, studies have reported the existence of CRISPR-Cas systems in phages, confering a survival advantage to the host organism (ref^27^ Seed et al., 2013). Interestingly, 33% of these viral contigs were also found within a metagenome-assembled genome (MAG) constructed from the same dataset in our earlier study (ref^54^ Duru et al., 2023), suggesting the possibility that the cas defense systems in these MAGs may have been acquired from the phages. Finally, we have noted the presence of several CRISPR arrays within the viral contigs, lacking any cas genes, with the majority consisting of less than three repeats. In bacterial genomes, it is customary to exclude CRISPR arrays containing less than three repeats. However, a prior study has demonstrated the existence of mini-CRISPR arrays in viruses that lack cas genes (ref^28^ Medvedeva et al., 2019). As a result, we have refrained from filtering out CRISPR arrays containing less than three repeats, resulting in the identification of 1 345 viral CRISPR arrays containing less than three repeats.

Recently, there have been reports of alternative genetic codes being utilized in phages, which may offer certain adaptive advantages (ref^51^ Borges et al., 2022, ref^68^ Peters et al., 2022). To investigate genetic code usage within the GPD and our predicted viral contigs, we assessed the coding density of the phages. The most common standard code in both the GPD and our viral contigs was the standard code (code 11). For alternative code usage, we observed dramatic increases in coding density. For example, for the GPD phages that prefer to use code 15, the average coding density was 91% with code 15, while with the standard code it was only 63%. It appears that applying an alternative genetic code during the annotation process can greatly enhance the structural gene annotation for certain phages. It is noteworthy that the second most common genetic code in the GPD was code 4, whereas in our viral contigs it was code 15. It should be emphasized that coding density calculation is more accurate when the phage is complete or nearly complete (ref^51^ Borges et al., 2022). We did not filter out non-complete phage contigs in our study, and most of the phages were categorized as “low quality draft”. This difference in genetic code preference distribution between GPD phages and our phages may arise for this reason.

Several studies have reported differences in the microbial composition of the gut microbiome in Parkinson’s disease patients relative to healthy controls (ref^69^ Boertien et al., 2019). Various strategies can be employed to manipulate the gut microbiome, such as prebiotics, environmental factors, and physical activity, but antibiotics and probiotics are particularly effective in altering the microbial population structure in the gut. In addition, several studies have shown that the use of a combination of bacteriophages, known as a “cocktail of phage”, can effectively modify the structural composition of the gut microbial community (ref^70^ Febvre et al., 2019, ref^71^ Federici et al., 2022, ref^72^ Galtier et al., 2017). In summary, this study provides evidence of changes in mobile genetic element populations in the gut microbiome of Parkinson’s disease patients. It is possible that phages could serve as biomarkers or even a treatment option if suitable phages are identified and isolated. However, this would require significant effort as identifying and isolating appropriate phages is a complex process.

## Acknowledgments

The authors thank the personnel of the DNA Sequencing and Genomics Laboratory for running the NGS assays. We acknowledge the CSC - IT Center for Science, Finland, for computational resources, and the University of Helsinki Language Services for English language revision.

## Author contributions

AL., P.A.B.P., and P.A. conceived and designed the study. F.S. performed clinical evaluation of the patients. P.A.B.P provided statistical support. I.C.D, A.L., T.K.S., P.L., and J.S. analyzed the sequencing data. L.P organized NGS assays. AL., drafted the manuscript. All authors reviewed and approved the final manuscript.

## Data availability

All sequencing data have been deposited in the European Nucleotide Archive (ENA) under accession code PRJEB59350.

## Notes

### Competing Interest Statement

FS received grants from the Academy of Finland, the Hospital District of Helsinki and Uusimaa, the OLVI Foundation, Konung Gustaf V:s och Drottning Victorias Frimurarestiftelse, the Wilhelm and Else Stockmann Foundation, the Emil Aaltonen Foundation, the Yrjo Jahnsson Foundation and the Sigrid Juselius Foundation, Renishaw. Honoraria: AbbVie, Axial Biotherapeutics, Orion, GE Healthcare, Merck, Teva, Bristol Myers Squibb, Sanofi, Biocodex, Lundbeck, and Biogen. FS is th founder and CEO of NeuroInnovation Oy and NeuroBiome Ltd. and is a member of the advisory boards of Axial Biotherapeutics and MRM Health. He has stock options from Axial Biotherapeutics.
P.A.B.P., L.P., P.A., and F.S. have patents issued (FI127671B, US10139408B2, US11499971B2) and pending (US16/186,663, EP3149205) that are assigned to NeuroBiome Ltd.
T.K.S. was funded by the Novo Nordisk Foundation (NNF22OC0080109).

## References

1. Dorsey, E. R. et al. Projected number of people with Parkinson disease in the most populous nations, 2005 through 2030. Neurology 68, 384–386 (2007).

2. Scheperjans, F. et al. Gut microbiota are related to Parkinson’s disease and clinical phenotype. Mov. Disord. Off. J. Mov. Disord. Soc. 30, 350–358 (2015).

3. Braak, H. et al. Staging of brain pathology related to sporadic Parkinson’s disease. Neurobiol. Aging 24, 197–211 (2003).

4. Braak, H., de Vos, R. A. I., Bohl, J. & Del Tredici, K. Gastric alpha-synuclein immunoreactive inclusions in Meissner’s and Auerbach’s plexuses in cases staged for Parkinson’s disease-related brain pathology. Neurosci. Lett. 396, 67–72 (2006).

5. Cersosimo, M. G. & Benarroch, E. E. Pathological correlates of gastrointestinal dysfunction in Parkinson’s disease. Neurobiol. Dis. 46, 559–564 (2012).

6. Mertsalmi, T. H. et al. More than constipation - bowel symptoms in Parkinson’s disease and their connection to gut microbiota. Eur. J. Neurol. 24, 1375–1383 (2017).

7. Perez-Pardo, P. et al. The gut-brain axis in Parkinson’s disease: Possibilities for food-based therapies. Eur. J. Pharmacol. 817, 86–95 (2017).

8. Wang, H.-X. & Wang, Y.-P. Gut Microbiota-brain Axis. Chin. Med. J. (Engl.) 129, 2373–2380 (2016).

9. Iannone, L. F. et al. Microbiota-gut brain axis involvement in neuropsychiatric disorders. Expert Rev. Neurother. 19, 1037–1050 (2019).

10. Cryan, J. F., O’Riordan, K. J., Sandhu, K., Peterson, V. & Dinan, T. G. The gut microbiome in neurological disorders. Lancet Neurol. 19, 179–194 (2020).

11. Blander, J. M., Longman, R. S., Iliev, I. D., Sonnenberg, G. F. & Artis, D. Regulation of inflammation by microbiota interactions with the host. Nat. Immunol. 18, 851–860 (2017).

12. Aho, V. T. E. et al. Relationships of gut microbiota, short-chain fatty acids, inflammation, and the gut barrier in Parkinson’s disease. Mol. Neurodegener. 16, 6 (2021).

13. Cryan, J. F. & Dinan, T. G. Mind-altering microorganisms: the impact of the gut microbiota on brain and behaviour. Nat. Rev. Neurosci. 13, 701–712 (2012).

14. Bonaz, B., Bazin, T. & Pellissier, S. The Vagus Nerve at the Interface of the Microbiota-Gut-Brain Axis. Front. Neurosci. 12, 49 (2018).

15. Aho, V. T. E. et al. Gut microbiota in Parkinson’s disease: Temporal stability and relations to disease progression. EBioMedicine 44, 691–707 (2019).

16. Breitbart, M. et al. Metagenomic analyses of an uncultured viral community from human feces. J. Bacteriol. 185, 6220–6223 (2003).

17. Minot, S. et al. The human gut virome: inter-individual variation and dynamic response to diet. Genome Res. 21, 1616–1625 (2011).

18. Minot, S., Grunberg, S., Wu, G. D., Lewis, J. D. & Bushman, F. D. Hypervariable loci in the human gut virome. Proc. Natl. Acad. Sci. U. S. A. 109, 3962–3966 (2012).

19. Minot, S. et al. Rapid evolution of the human gut virome. Proc. Natl. Acad. Sci. 110, 12450–12455 (2013).

20. Hannigan, G. D., Duhaime, M. B., Ruffin, M. T., Koumpouras, C. C. & Schloss, P. D. Diagnostic Potential and Interactive Dynamics of the Colorectal Cancer Virome. mBio 9, e02248–18 (2018).

21. Yang, K. et al. Alterations in the Gut Virome in Obesity and Type 2 Diabetes Mellitus. Gastroenterology 161, 1257–1269.e13 (2021).

22. Liang, G., Cobián-Güemes, A. G., Albenberg, L. & Bushman, F. The gut virome in inflammatory bowel diseases. Curr. Opin. Virol. 51, 190–198 (2021).

23. Adiliaghdam, F., et al. Human enteric viruses autonomously shape inflammatory bowel disease phenotype through divergent innate immunomodulation. Sci. Immunol. 7, eabn6660 (2022).

24. Draper, L. A. et al. Autochthonous faecal viral transfer (FVT) impacts the murine microbiome after antibiotic perturbation. BMC Biol. 18, 173 (2020).

25. Camarillo-Guerrero, L. F., Almeida, A., Rangel-Pineros, G., Finn, R. D. & Lawley, T. D. Massive expansion of human gut bacteriophage diversity. Cell 184, 1098–1109.e9 (2021).

26. Hille, F. & Charpentier, E. CRISPR-Cas: biology, mechanisms and relevance. Philos. Trans. R. Soc. B Biol. Sci. 371, 20150496 (2016).

27. Seed, K. D., Lazinski, D. W., Calderwood, S. B. & Camilli, A. A bacteriophage encodes its own CRISPR/Cas adaptive response to evade host innate immunity. Nature 494, 489–491 (2013).

28. Medvedeva, S. et al. Virus-borne mini-CRISPR arrays are involved in interviral conflicts. Nat. Commun. 10, 5204 (2019).

29. Bedarf, J. R. et al. Functional implications of microbial and viral gut metagenome changes in early stage L-DOPA-naïve Parkinson’s disease patients. Genome Med. 9, 39 (2017).

30. Wallen, Z. D. et al. Metagenomics of Parkinson’s disease implicates the gut microbiome in multiple disease mechanisms. Nat. Commun. 13, 6958 (2022).

31. Qian, Y. et al. Gut metagenomics-derived genes as potential biomarkers of Parkinson’s disease. Brain J. Neurol. 143, 2474–2489 (2020).

32. Berardelli, A. et al. EFNS/MDS-ES/ENS [corrected] recommendations for the diagnosis of Parkinson’s disease. Eur. J. Neurol. 20, 16–34 (2013).

33. Martin, M. Cutadapt removes adapter sequences from high-throughput sequencing reads. EMBnet.journal 17, 10–12 (2011).

34. Li, H. & Durbin, R. Fast and accurate short read alignment with Burrows-Wheeler transform. Bioinforma. Oxf. Engl. 25, 1754–1760 (2009).

35. Nurk, S., Meleshko, D., Korobeynikov, A. & Pevzner, P. A. metaSPAdes: a new versatile metagenomic assembler. Genome Res. 27, 824 (2017).

36. Nurk, S. et al. Assembling Genomes and Mini-metagenomes from Highly Chimeric Reads. in Research in Computational Molecular Biology (eds. Deng, M., Jiang, R., Sun, F. & Zhang, X.) 158–170 (Springer, 2013). doi:10.1007/978-3-642-37195-0_13.

37. Yu, M. K., Fogarty, E. C. & Eren, A. M. The genetic and ecological landscape of plasmids in the human gut. bioRxiv 2020.11.01.361691 (2022) doi:10.1101/2020.11.01.361691.

38. Seemann, T. Prokka: rapid prokaryotic genome annotation. Bioinforma. Oxf. Engl. 30, 2068–2069 (2014).

39. Sequeira, J. C., Rocha, M., Alves, M. M. & Salvador, A. F. UPIMAPI, reCOGnizer and KEGGCharter: Bioinformatics tools for functional annotation and visualization of (meta)-omics datasets. Comput. Struct. Biotechnol. J. 20, 1798–1810 (2022).

40. Menzel, P., Ng, K. L. & Krogh, A. Fast and sensitive taxonomic classification for metagenomics with Kaiju. Nat. Commun. 7, 11257 (2016).

41. Edgar, R. C. PILER-CR: Fast and accurate identification of CRISPR repeats. BMC Bioinformatics 8, 18 (2007).

42. Langmead, B. & Salzberg, S. L. Fast gapped-read alignment with Bowtie 2. Nat. Methods 9, 357–359 (2012).

43. R: The R Project for Statistical Computing. https://www.r-project.org/.

44. Oksanen, J., et al. vegan: Community Ecology Package. (2012).

45. Love, M. I., Huber, W. & Anders, S. Moderated estimation of fold change and dispersion for RNA-seq data with DESeq2. Genome Biol. 15, 550 (2014).

46. Kieft, K., Zhou, Z. & Anantharaman, K. VIBRANT: automated recovery, annotation and curation of microbial viruses, and evaluation of viral community function from genomic sequences. Microbiome 8, 90 (2020).

47. Russel, J., Pinilla-Redondo, R., Mayo-Muñoz, D., Shah, S. A. & Sørensen, S. J. CRISPRCasTyper: Automated Identification, Annotation, and Classification of CRISPR-Cas Loci. CRISPR J. 3, 462–469 (2020).

48. Zhang, R. et al. SpacePHARER: Sensitive identification of phages from CRISPR spacers in prokaryotic hosts. Bioinforma. Oxf. Engl. 37, 3364–3366 (2021).

49. Guerin, E. et al. Biology and Taxonomy of crAss-like Bacteriophages, the Most Abundant Virus in the Human Gut. Cell Host Microbe 24, 653–664.e6 (2018).

50. Camacho, C. et al. BLAST+: architecture and applications. BMC Bioinformatics 10, 421 (2009).

51. Borges, A. L. et al. Widespread stop-codon recoding in bacteriophages may regulate translation of lytic genes. Nat. Microbiol. 7, 918–927 (2022).

52. Pedregosa, F. et al. Scikit-learn: Machine Learning in Python. Mach. Learn. PYTHON.

53. Lundberg, S. M. et al. From local explanations to global understanding with explainable AI for trees. Nat. Mach. Intell. 2, 56–67 (2020).

54. Duru, I. C. et al. Metagenome-assembled microbial genomes from Parkinson’s disease fecal samples. bioRxiv 2023.02.27.526590 (2023) doi:10.1101/2023.02.27.526590.

55. Dutilh, B. E. et al. A highly abundant bacteriophage discovered in the unknown sequences of human faecal metagenomes. Nat. Commun. 5, 4498 (2014).

56. Parashar, A. & Udayabanu, M. Gut microbiota: Implications in Parkinson’s disease. Parkinsonism Relat. Disord. 38, 1–7 (2017).

57. Hopfner, F. et al. Gut microbiota in Parkinson disease in a northern German cohort. Brain Res. 1667, 41–45 (2017).

58. Mao, L. et al. Cross-Sectional Study on the Gut Microbiome of Parkinson’s Disease Patients in Central China. Front. Microbiol. 12, (2021).

59. Suzuki, Y. et al. Long-read metagenomic exploration of extrachromosomal mobile genetic elements in the human gut. Microbiome 7, 119 (2019).

60. Broaders, E., Gahan, C. G. M. & Marchesi, J. R. Mobile genetic elements of the human gastrointestinal tract: potential for spread of antibiotic resistance genes. Gut Microbes 4, 271–280 (2013).

61. Kurokawa, K. et al. Comparative Metagenomics Revealed Commonly Enriched Gene Sets in Human Gut Microbiomes. DNA Res. 14, 169–181 (2007).

62. Smillie, C. S. et al. Ecology drives a global network of gene exchange connecting the human microbiome. Nature 480, 241–244 (2011).

63. Liu, B.-N., Liu, X.-T., Liang, Z.-H. & Wang, J.-H. Gut microbiota in obesity. World J. Gastroenterol. 27, 3837–3850 (2021).

64. Odamaki, T. et al. Age-related changes in gut microbiota composition from newborn to centenarian: a cross-sectional study. BMC Microbiol. 16, 90 (2016).

65. Tisza, M. J. & Buck, C. B. A catalog of tens of thousands of viruses from human metagenomes reveals hidden associations with chronic diseases. Proc. Natl. Acad. Sci. 118, e2023202118 (2021).

66. Sausset, R., Petit, M. A., Gaboriau-Routhiau, V. & De Paepe, M. New insights into intestinal phages. Mucosal Immunol. 13, 205–215 (2020).

67. Bhardwaj, K. et al. Insights into the human gut virome by sampling a population from the Indian subcontinent. J. Gen. Virol. 103, (2022).

68. Peters, S. L. et al. Experimental validation that human microbiome phages use alternative genetic coding. Nat. Commun. 13, 5710 (2022).

