## Supplementary Fig. S6 for "Comparison of phage and plasmid populations present in the gut microbiota of Parkinson’s disease patients": Supplementary_Figure_S6.pdf

a)

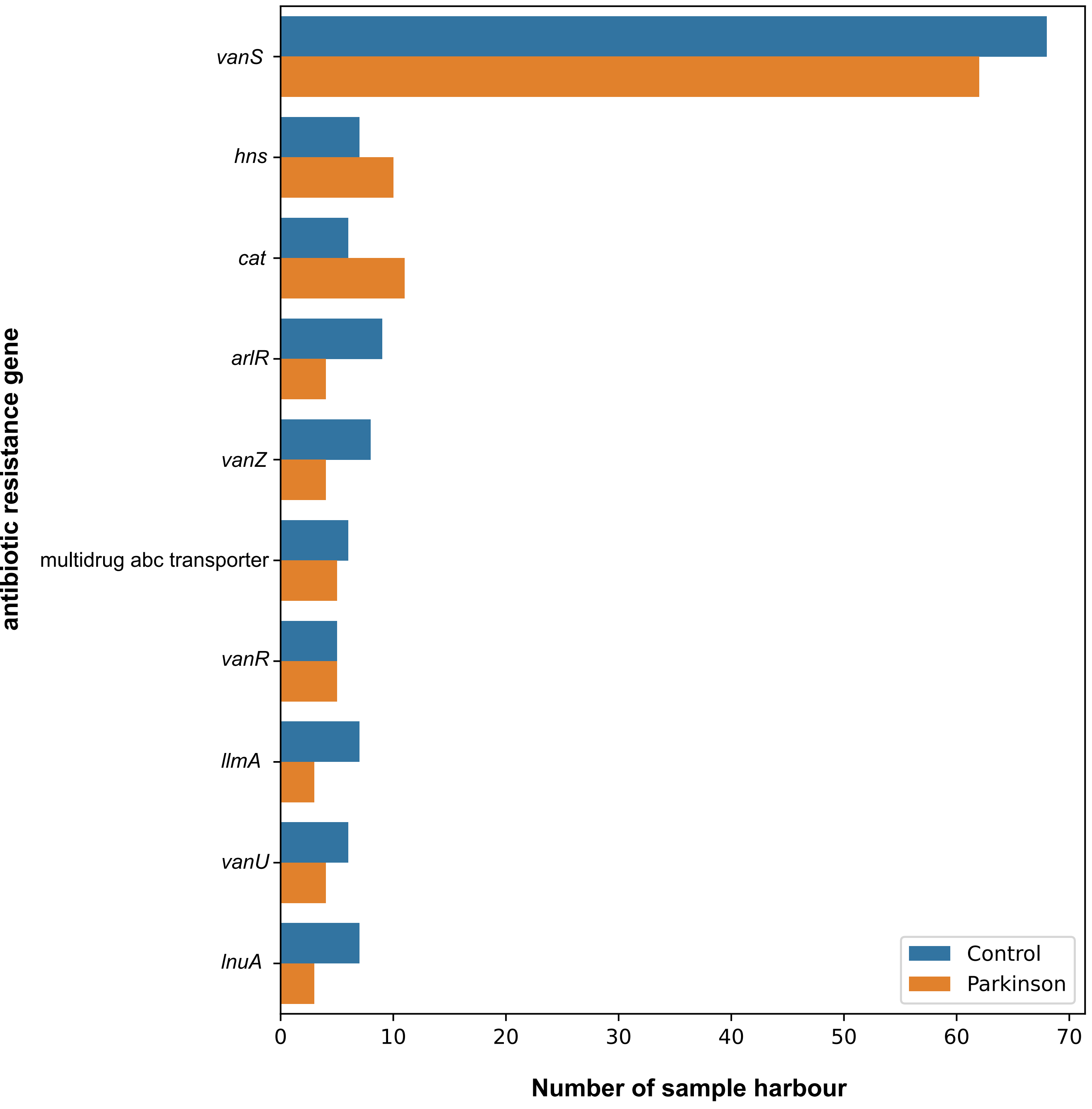


b)

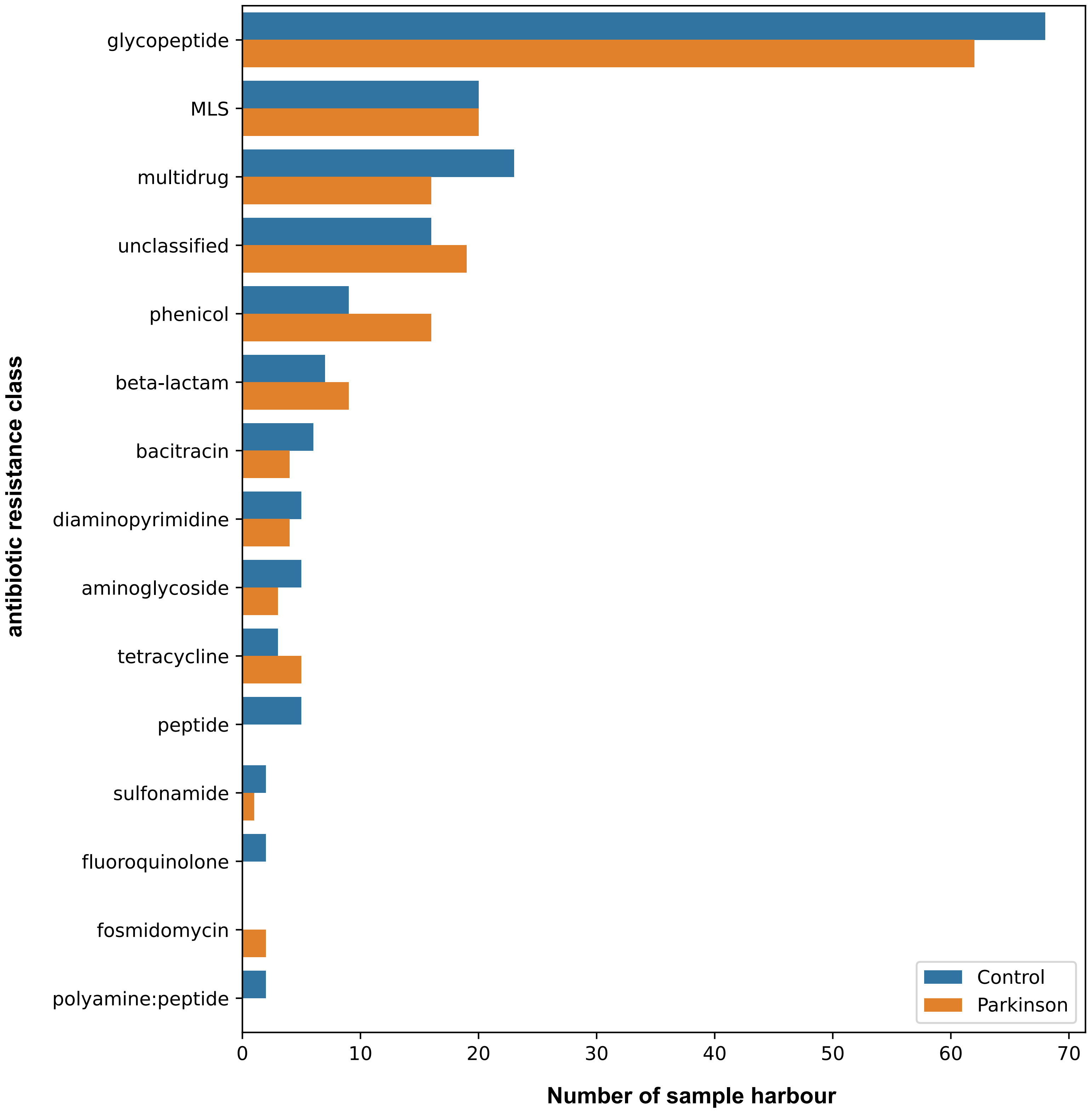


Supplementary Figure S6. The count plots of (a) antibiotic resistance genes and (b) antibiotic resistance classes within plasmid contigs. The blue bar represent the control samples, while the orange bars represent the PD samples.
