## Supplementary Fig. S9 for "Comparison of phage and plasmid populations present in the gut microbiota of Parkinson’s disease patients": Supplementary_Figure_S9.pdf

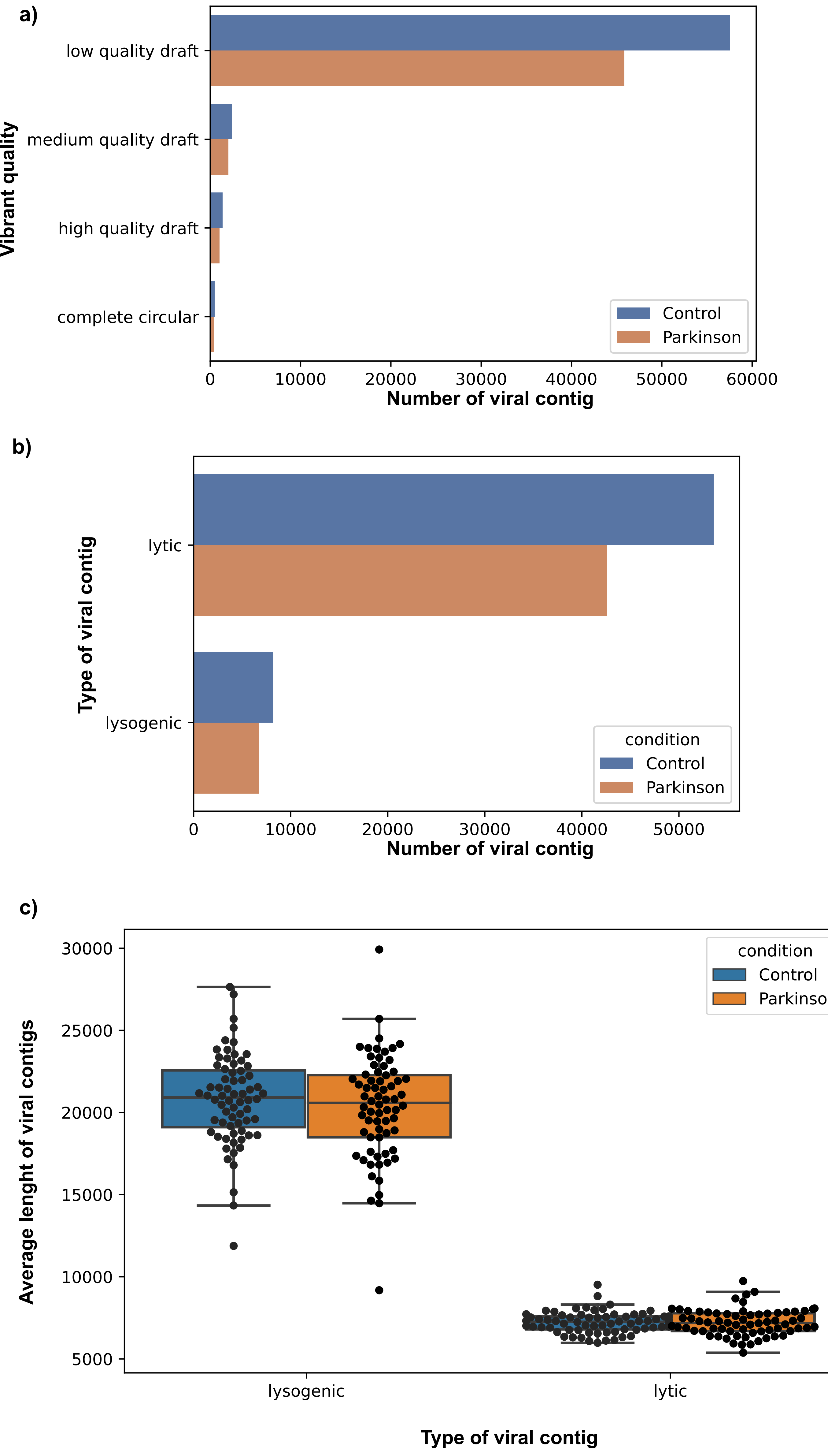


Supplementary Figure S9. a) Bar chart shows the number of predicted viral contigs and their quality classification based on Vibrant. b) Bar chart shows the number of lytic and lysogenic viral contigs. c) The average length of viral contigs are shown using box plots. Each dot represents one sample. Blue bar represents the control group, orange represents the Parkinson group.
