## Supplementary Fig. S5 for "Comparison of phage and plasmid populations present in the gut microbiota of Parkinson’s disease patients": Supplementary_Figure_S5.pdf

a)

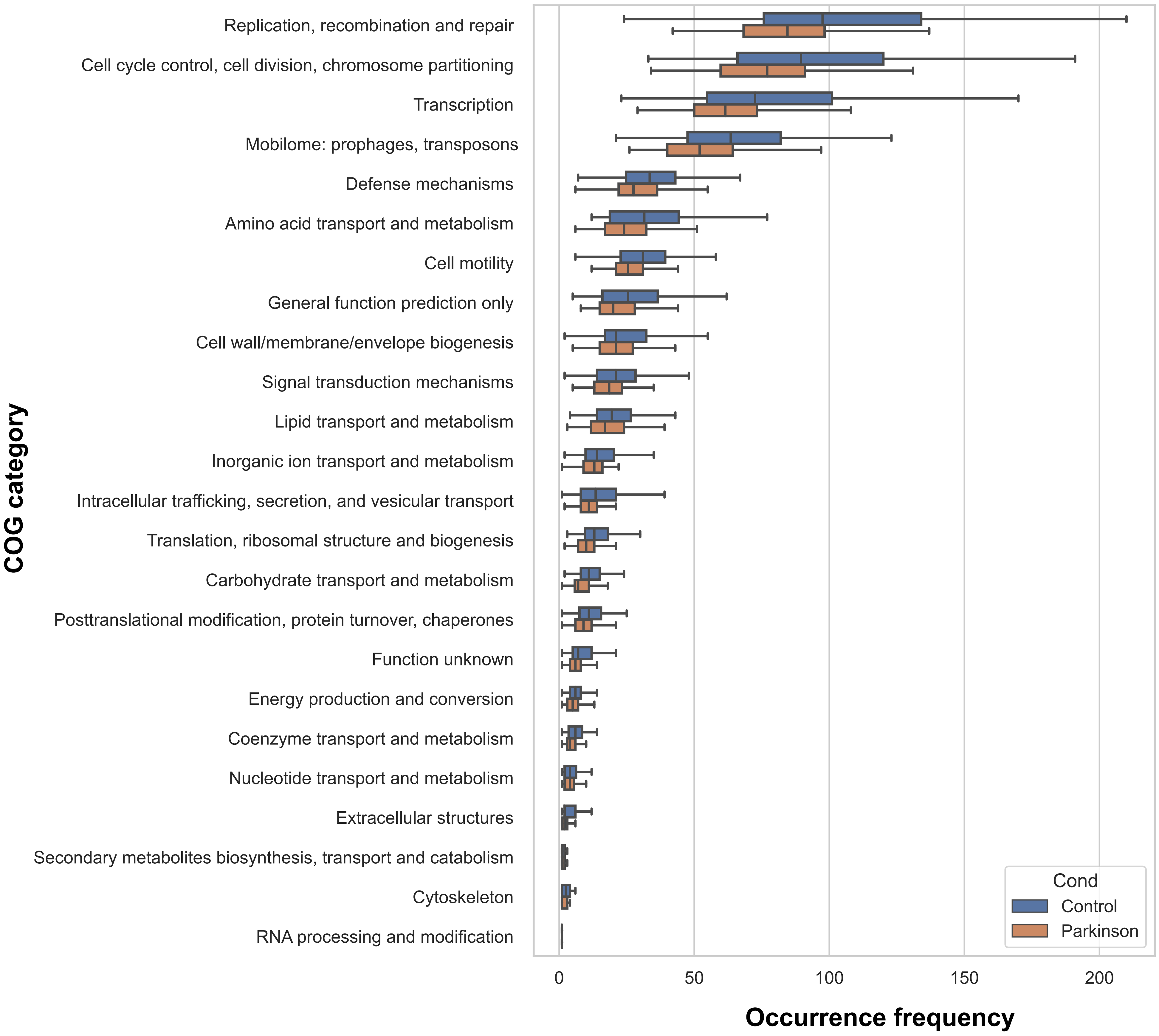


b)

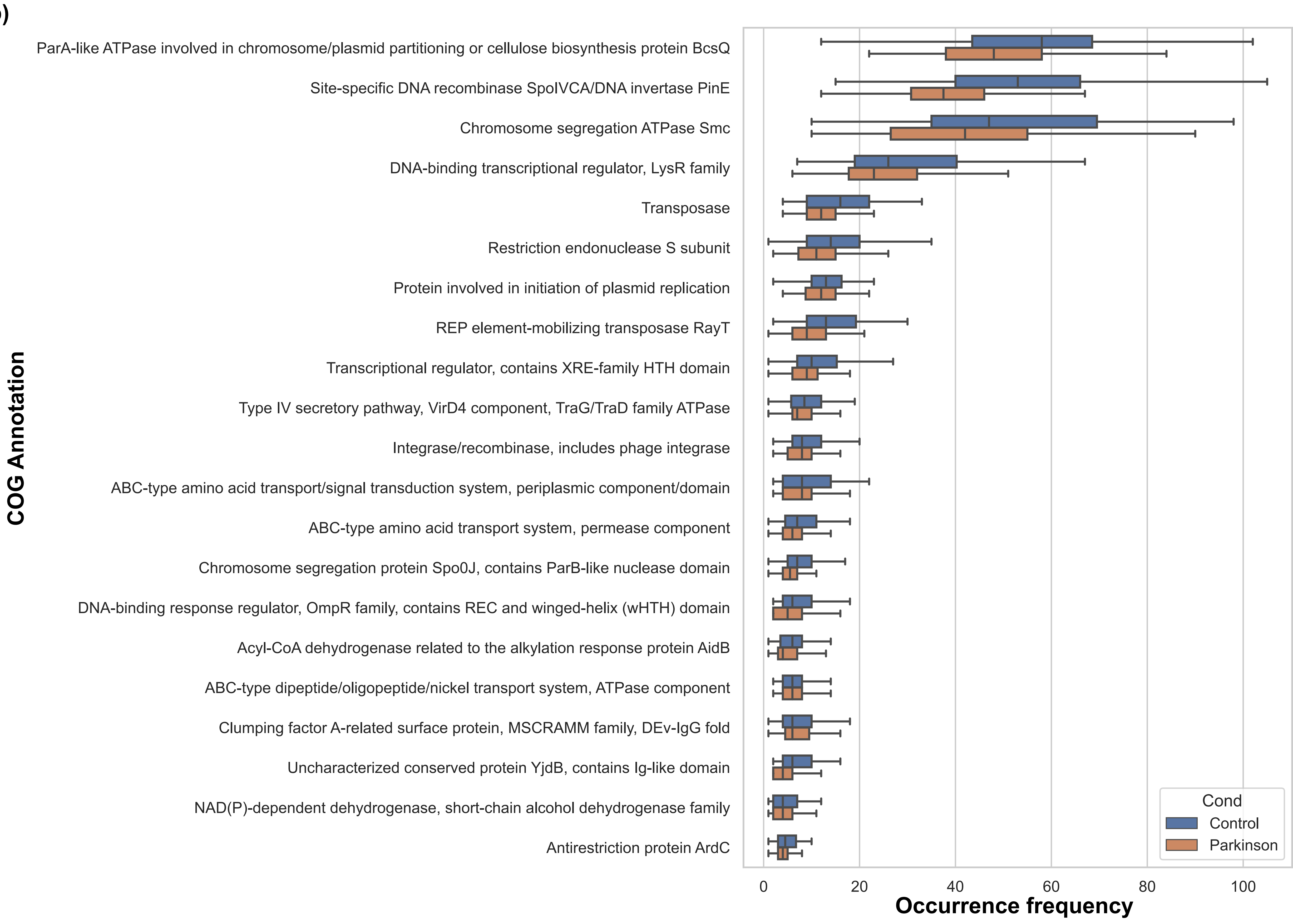


Supplementary Figure S5. The occurence frequency of (a) COG category and (b) COG annotation for plasmid genes. The box plot summarise the distribution of occurence frequencies with a vertical line indicating the median, and a rectangular box that spans interquartile range. Blue box represents control samples, and orange represent PD samples.
