## Supplementary Fig. S10 for "Comparison of phage and plasmid populations present in the gut microbiota of Parkinson’s disease patients": Supplementary_Figure_S10.pdf

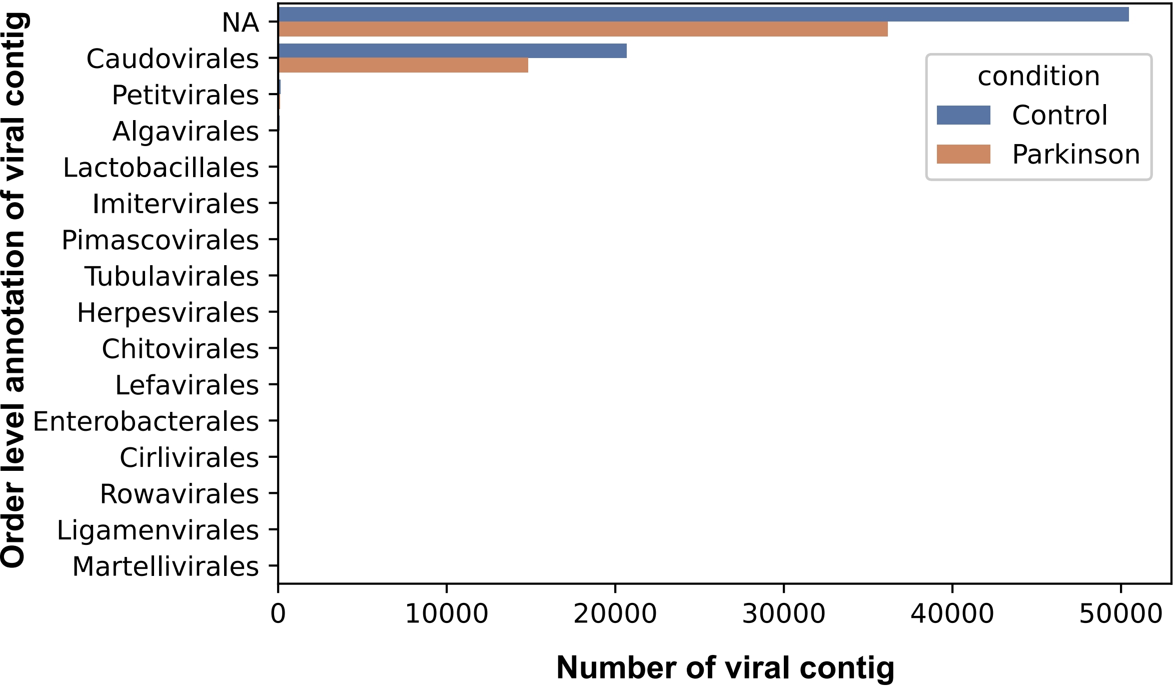


| Order | #contig control | #contig parkinson |
|---|---|---|
| NA | 50468 | 36165 |
| Caudovirales | 20668 | 14813 |
| Petitvirales | 134 | 110 |
| Algavirales | 62 | 47 |
| Lactobacillales | 35 | 27 |
| Imitervirales | 24 | 27 |
| Pimascovirales | 32 | 13 |
| Herpesvirales | 7 | 4 |
| Tubulavirales | 8 | 4 |
| Enterobacterales | 1 | 3 |
| Lefavirales | 2 | 2 |
| Cirlivirales | 1 | 1 |
| Chitovirales | 6 | 0 |
| Rowavirales | 2 | 0 |
| Ligamenvirales | 1 | 0 |
| Martellivirales | 1 | 0 |

Supplementary Figure S10. Bar chart shows the number of viral contigs and their order level classification. The table shows exact numbers, since small numbers cannot be seen in the bar chart. Blue bar represents the control group, yellow represents the PD group.
