## Supplementary Fig. S4 for "Comparison of phage and plasmid populations present in the gut microbiota of Parkinson’s disease patients": Supplementary_Figure_S4.pdf

**a) plasmid**

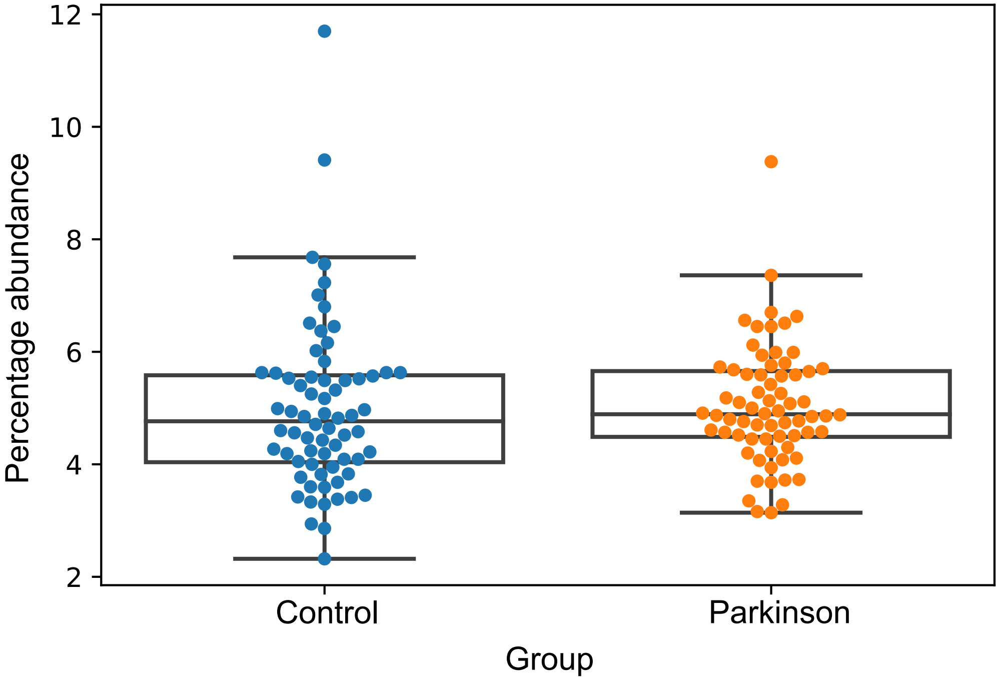


**b) phage**

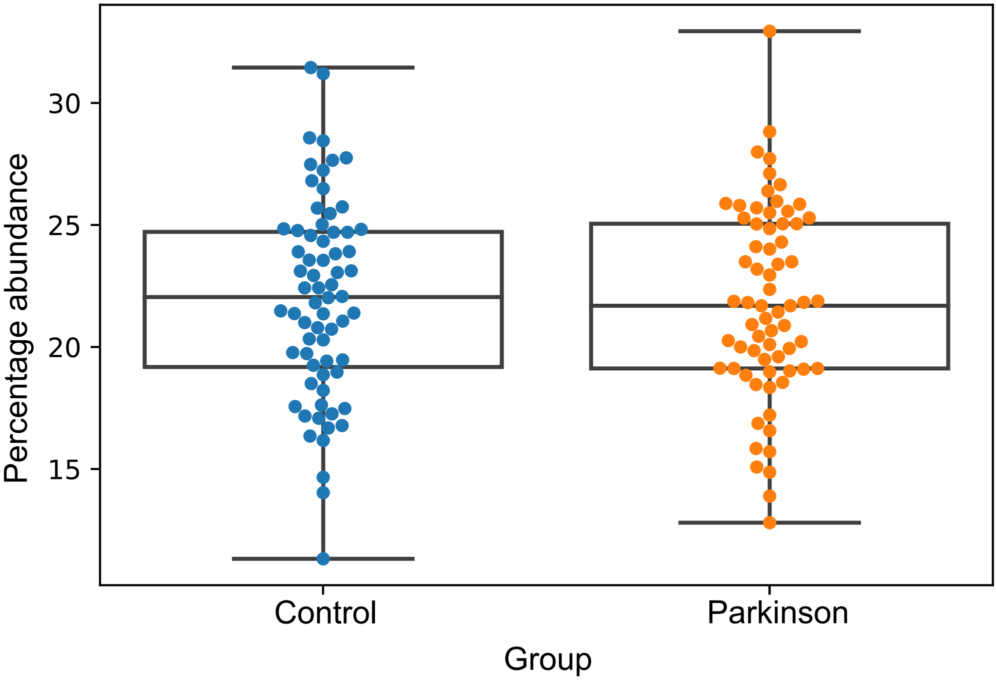


Supplementary Figure S4. The total (a) plasmid and (b) phage abundance per sample. The box plot summarise the distribution of sample abundances with a horizontal line indicating the median, and a rectangular box that spans interquartile range. Each sample also represented by a dot in the figure. Blue dot represents control samples, and orange represent PD samples.
