## Supplementary Fig. S2 for "Comparison of phage and plasmid populations present in the gut microbiota of Parkinson’s disease patients": Supplementary_Figure_S2.pdf

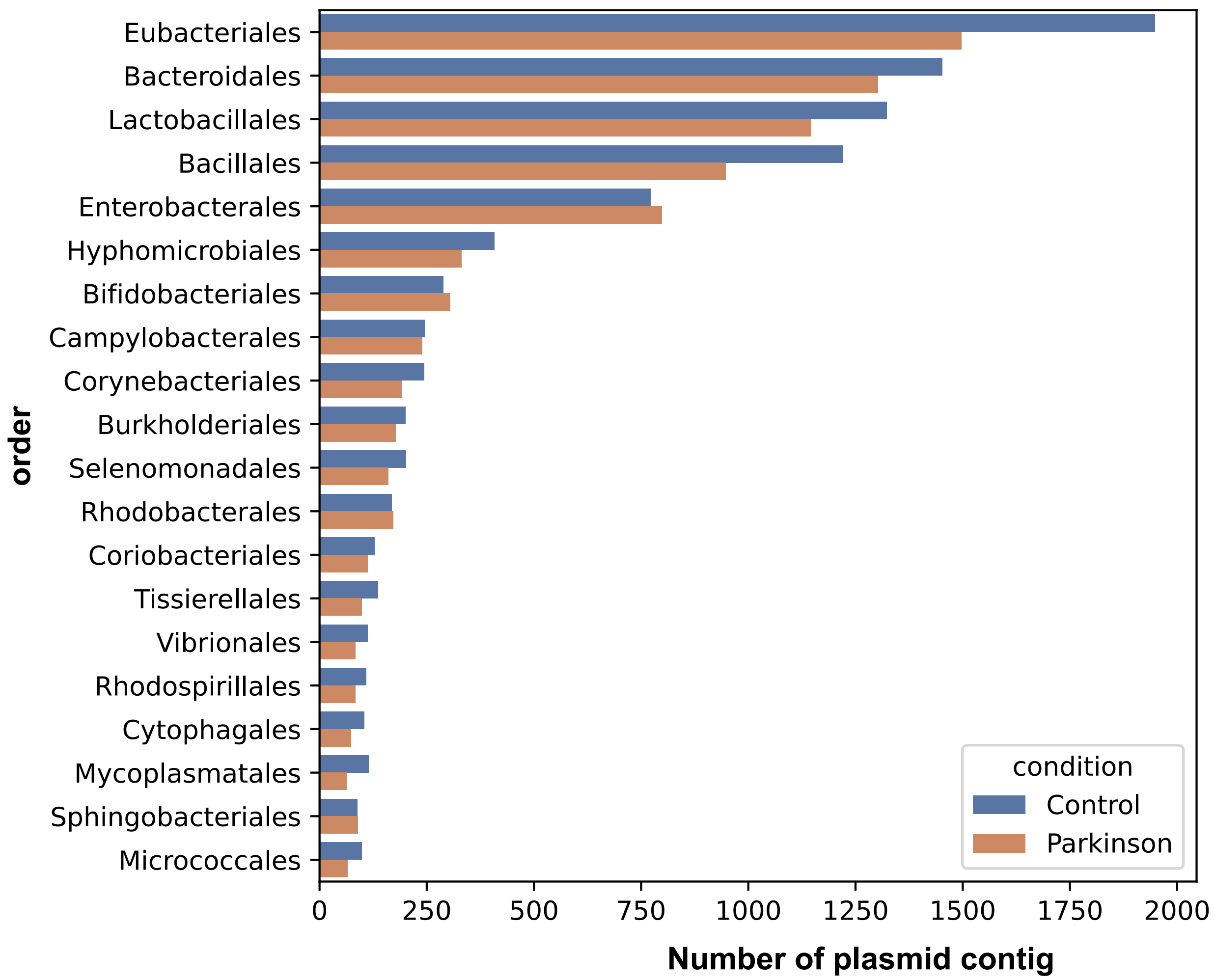
Eubacteriales
Bacteroidales
Lactobacillales
Bacillales
Enterobacterales
Hyphomicrobiales
Bifidobacteriales
Campylobacterales
Corynebacteriales
Burkholderiales
Selenomonadales
Rhodobacterales
Coriobacteriales
Tissierellales
Vibrionales
Rhodospirillales
Cytophagales
Mycoplasmatales
Sphingobacteriales
Micrococcales
order
condition
Control
Parkinson
0
250
500
750
1000
1250
1500
1750
2000
Number of plasmid contig

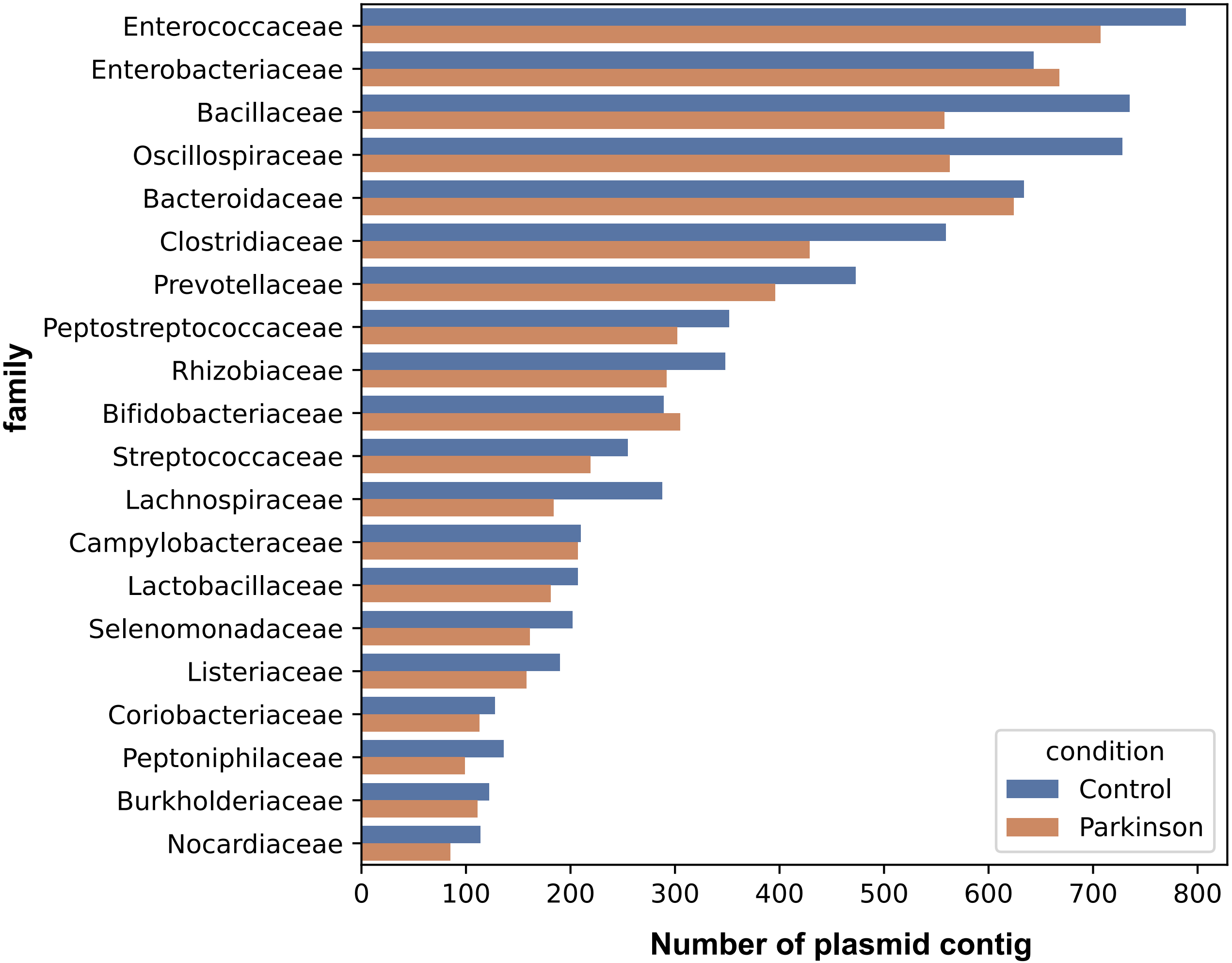
Enterococcaceae
Enterobacteriaceae
Bacillaceae
Oscillospiraceae
Bacteroidaceae
Clostridiaceae
Prevotellaceae
Peptostreptococcaceae
Rhizobiaceae
Bifidobacteriaceae
Streptococcaceae
Lachnospiraceae
Campylobacteraceae
Lactobacillaceae
Selenomonadaceae
Listeriaceae
Coriobacteriaceae
Peptoniphilaceae
Burkholderiaceae
Nocardiaceae
family
0
100
200
300
400
500
600
700
800
Number of plasmid contig
condition
Control
Parkinson

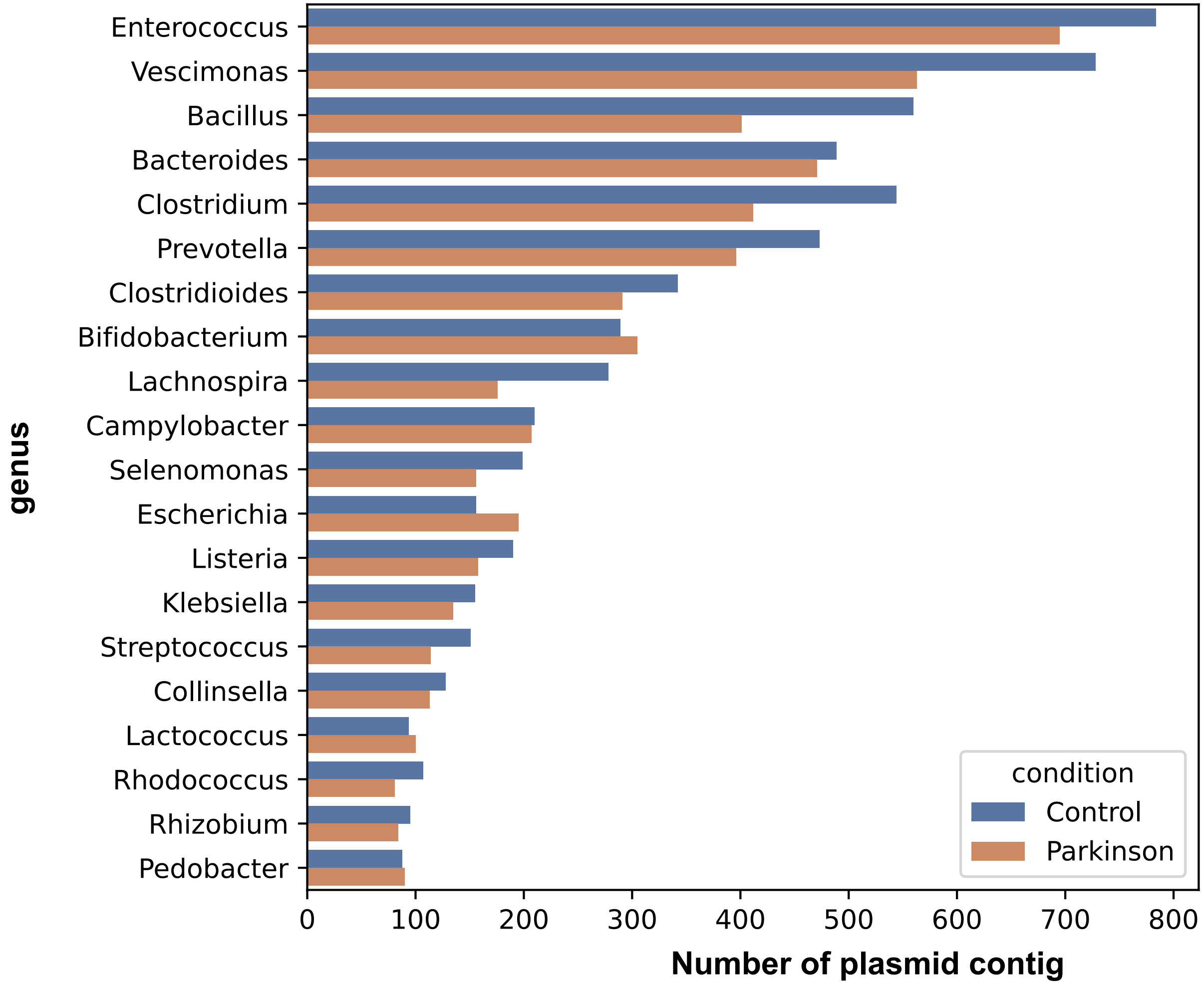
Enterococcus
Vescimonas
Bacillus
Bacteroides
Clostridium
Prevotella
Clostridioides
Bifidobacterium
Lachnospira
Campylobacter
Selenomonas
Escherichia
Listeria
Klebsiella
Streptococcus
Collinsella
Lactococcus
Rhodococcus
Rhizobium
Pedobacter
genus
0
100
200
300
400
500
600
700
800
Number of plasmid contig
condition
Control
Parkinson

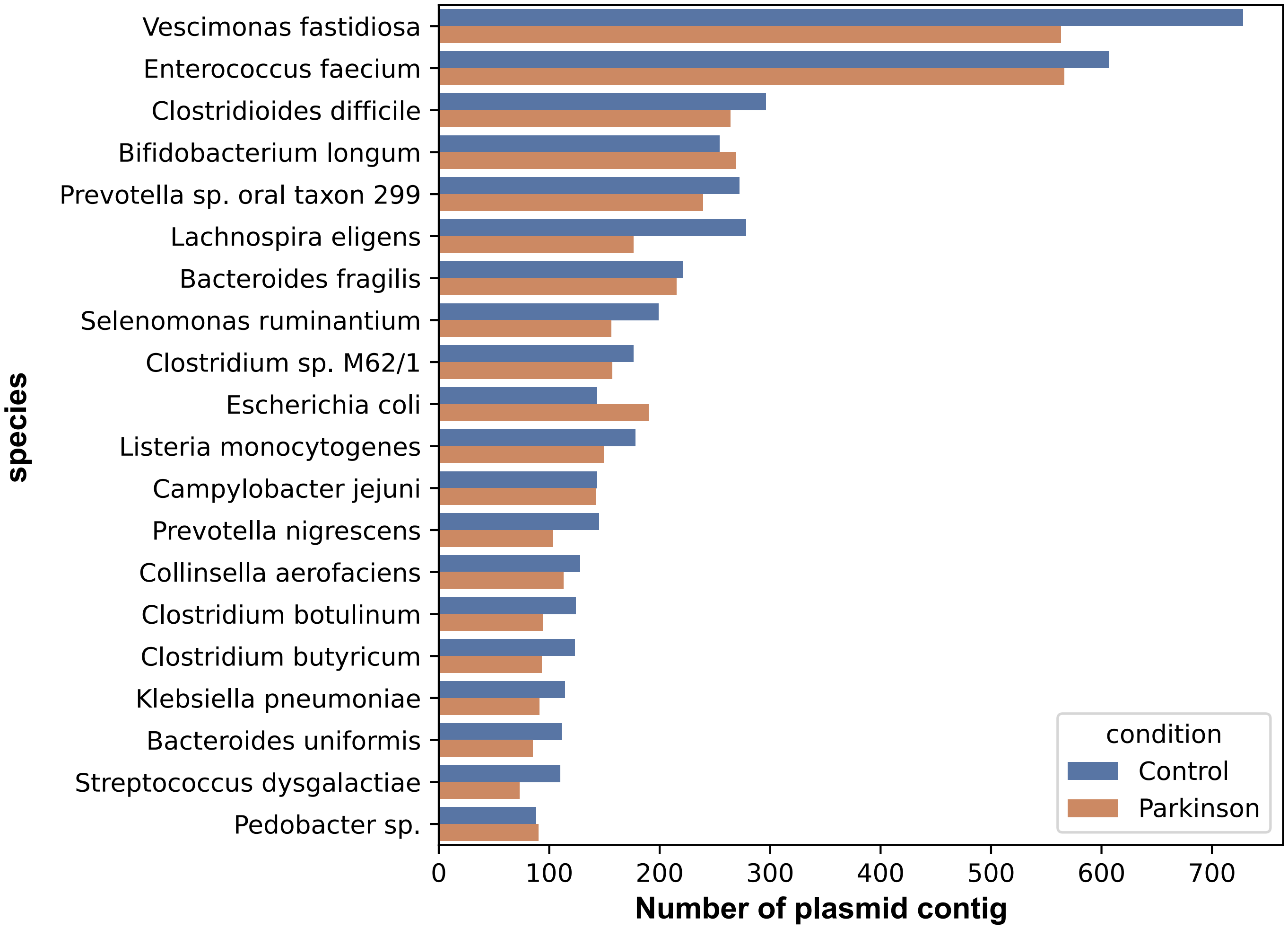
Vescimonas fastidiosa
Enterococcus faecium
Clostridioides difficile
Bifidobacterium longum
Prevotella sp. oral taxon 299
Lachnospira eligens
Bacteroides fragilis
Selenomonas ruminantium
Clostridium sp. M62/1
Escherichia coli
Listeria monocytogenes
Campylobacter jejuni
Prevotella nigrescens
Collinsella aerofaciens
Clostridium botulinum
Clostridium butyricum
Klebsiella pneumoniae
Bacteroides uniformis
Streptococcus dysgalactiae
Pedobacter sp.
species
0
100
200
300
400
500
600
700
Number of plasmid contig
condition
Control
Parkinson

Supplementary Figure S2. The countplots of annotations for every plasmid contig at different taxonomic levels, including order, family, genus, and species. The blue bars represent the control plasmid contigs, while the orange bars represent the PD plasmid contigs. To keep the figures concise, only the top 20 taxa are displayed..
