## Supplementary figures and images for "Comparison of phage and plasmid populations present in the gut microbiota of Parkinson’s disease patients"

### Supplementary_Figure_S1.pdf

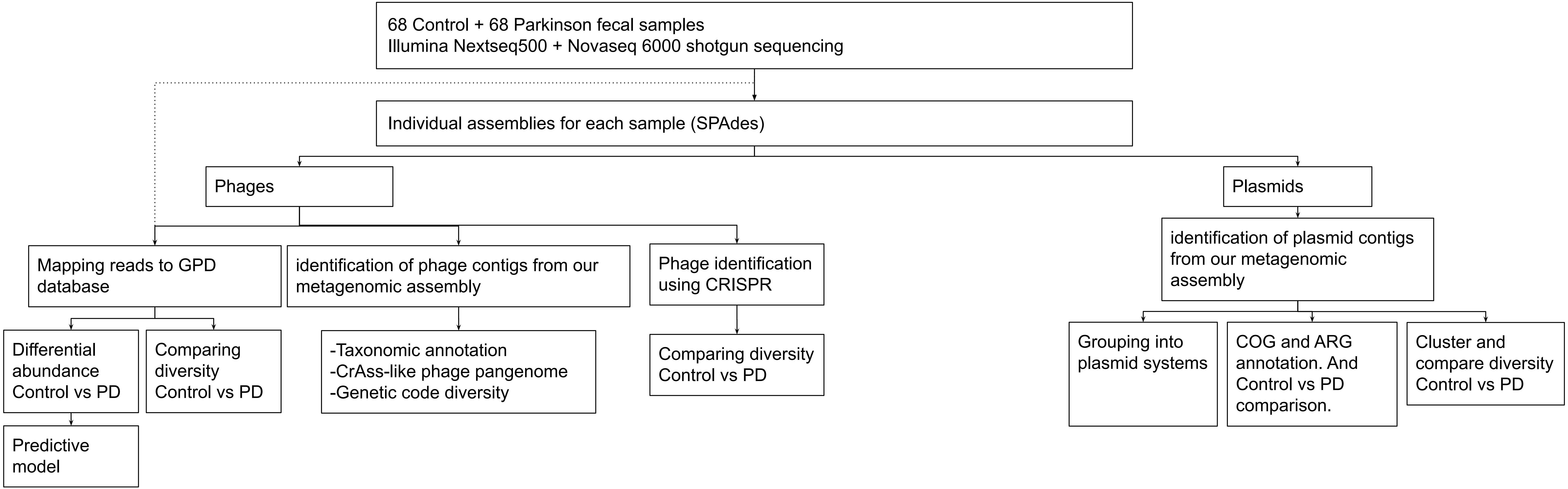


Supplementary Figure 1. A short summary of the pipeline that used in this study.
