## Supplementary Fig. S3 for "Comparison of phage and plasmid populations present in the gut microbiota of Parkinson’s disease patients": Supplementary Fig S3 Plasmid and Plasmid gene beta div plot.pptx

### Slide 1
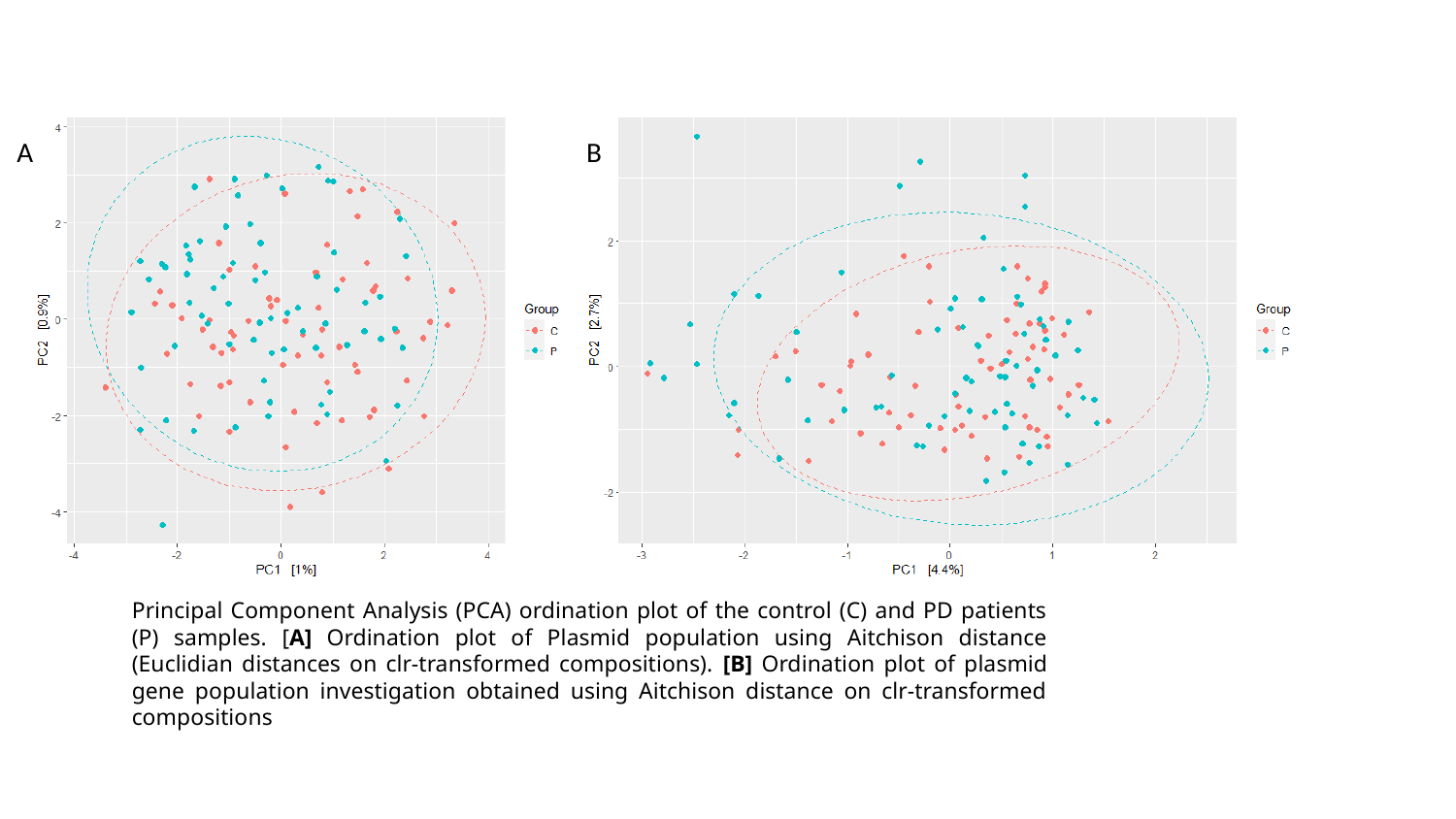

A
B
Principal Component Analysis (PCA) ordination plot of the control (C) and PD patients (P) samples. [A] Ordination plot of Plasmid population using Aitchison distance (Euclidian distances on clr-transformed compositions). [B] Ordination plot of plasmid gene population investigation obtained using Aitchison distance on clr-transformed compositions
