## Supplementary Fig. S8 for "Comparison of phage and plasmid populations present in the gut microbiota of Parkinson’s disease patients": Supplementary Figure S8 - ARG class.pptx

### Slide 1
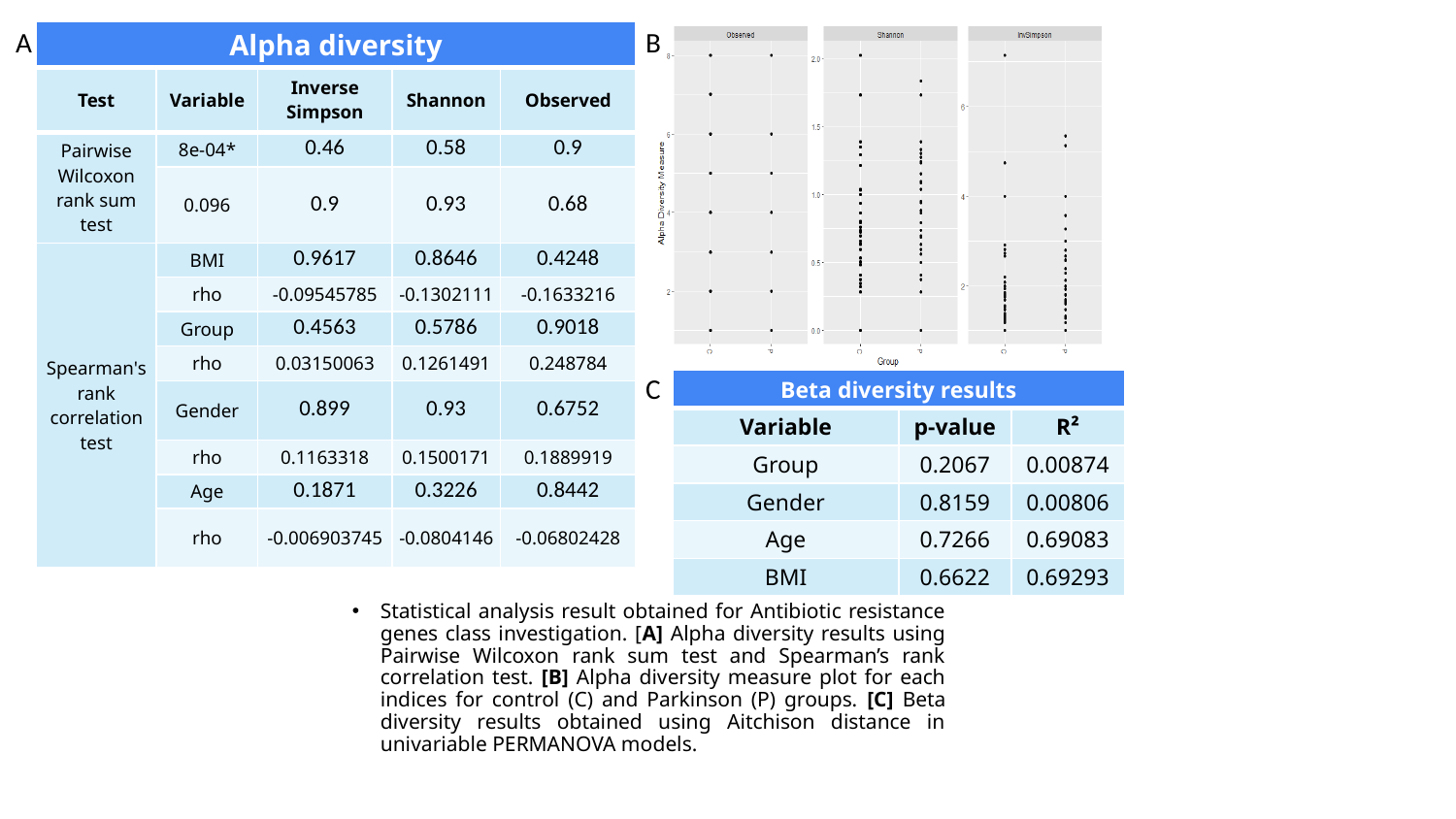

A
B
| Alpha diversity | | | | |
| --- | --- | --- | --- | --- |
| Test | Variable | Inverse Simpson | Shannon | Observed |
| Pairwise Wilcoxon rank sum test | 8e-04\* | 0.46 | 0.58 | 0.9 |
| | 0.096 | 0.9 | 0.93 | 0.68 |
| Spearman's rank correlation test | BMI | 0.9617 | 0.8646 | 0.4248 |
| | rho | -0.09545785 | -0.1302111 | -0.1633216 |
| | Group | 0.4563 | 0.5786 | 0.9018 |
| | rho | 0.03150063 | 0.1261491 | 0.248784 |
| | Gender | 0.899 | 0.93 | 0.6752 |
| | rho | 0.1163318 | 0.1500171 | 0.1889919 |
| | Age | 0.1871 | 0.3226 | 0.8442 |
| | rho | -0.006903745 | -0.0804146 | -0.06802428 |
C
| Beta diversity results | | |
| --- | --- | --- |
| Variable | p-value | R² |
| Group | 0.2067 | 0.00874 |
| Gender | 0.8159 | 0.00806 |
| Age | 0.7266 | 0.69083 |
| BMI | 0.6622 | 0.69293 |
Statistical analysis result obtained for Antibiotic resistance genes class investigation. [A] Alpha diversity results using Pairwise Wilcoxon rank sum test and Spearman’s rank correlation test. [B] Alpha diversity measure plot for each indices for control (C) and Parkinson (P) groups. [C] Beta diversity results obtained using Aitchison distance in univariable PERMANOVA models.
