## Supplementary Fig. S7 for "Comparison of phage and plasmid populations present in the gut microbiota of Parkinson’s disease patients": Supplementary Figure S7 - ARG.pptx

### Slide 1
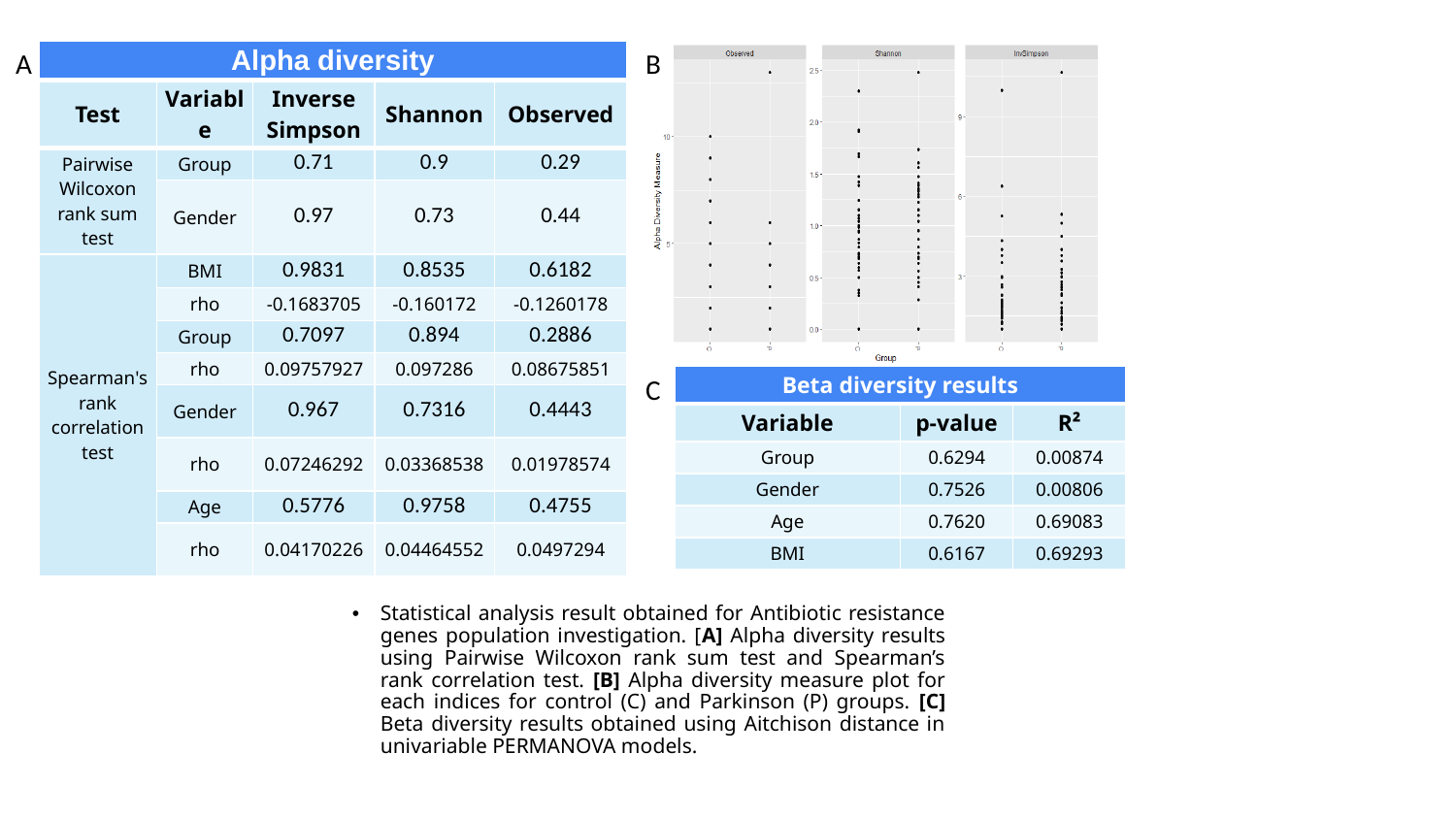

A
B
| Alpha diversity | | | | |
| --- | --- | --- | --- | --- |
| Test | Variable | Inverse Simpson | Shannon | Observed |
| Pairwise Wilcoxon rank sum test | Group | 0.71 | 0.9 | 0.29 |
| | Gender | 0.97 | 0.73 | 0.44 |
| Spearman's rank correlation test | BMI | 0.9831 | 0.8535 | 0.6182 |
| | rho | -0.1683705 | -0.160172 | -0.1260178 |
| | Group | 0.7097 | 0.894 | 0.2886 |
| | rho | 0.09757927 | 0.097286 | 0.08675851 |
| | Gender | 0.967 | 0.7316 | 0.4443 |
| | rho | 0.07246292 | 0.03368538 | 0.01978574 |
| | Age | 0.5776 | 0.9758 | 0.4755 |
| | rho | 0.04170226 | 0.04464552 | 0.0497294 |
C
| Beta diversity results | | |
| --- | --- | --- |
| Variable | p-value | R² |
| Group | 0.6294 | 0.00874 |
| Gender | 0.7526 | 0.00806 |
| Age | 0.7620 | 0.69083 |
| BMI | 0.6167 | 0.69293 |
Statistical analysis result obtained for Antibiotic resistance genes population investigation. [A] Alpha diversity results using Pairwise Wilcoxon rank sum test and Spearman’s rank correlation test. [B] Alpha diversity measure plot for each indices for control (C) and Parkinson (P) groups. [C] Beta diversity results obtained using Aitchison distance in univariable PERMANOVA models.
