## Supplementary Table S12 for "Comparison of phage and plasmid populations present in the gut microbiota of Parkinson’s disease patients": Supplementary table S12 - GPD lowest p-value.pptx

### Slide 1
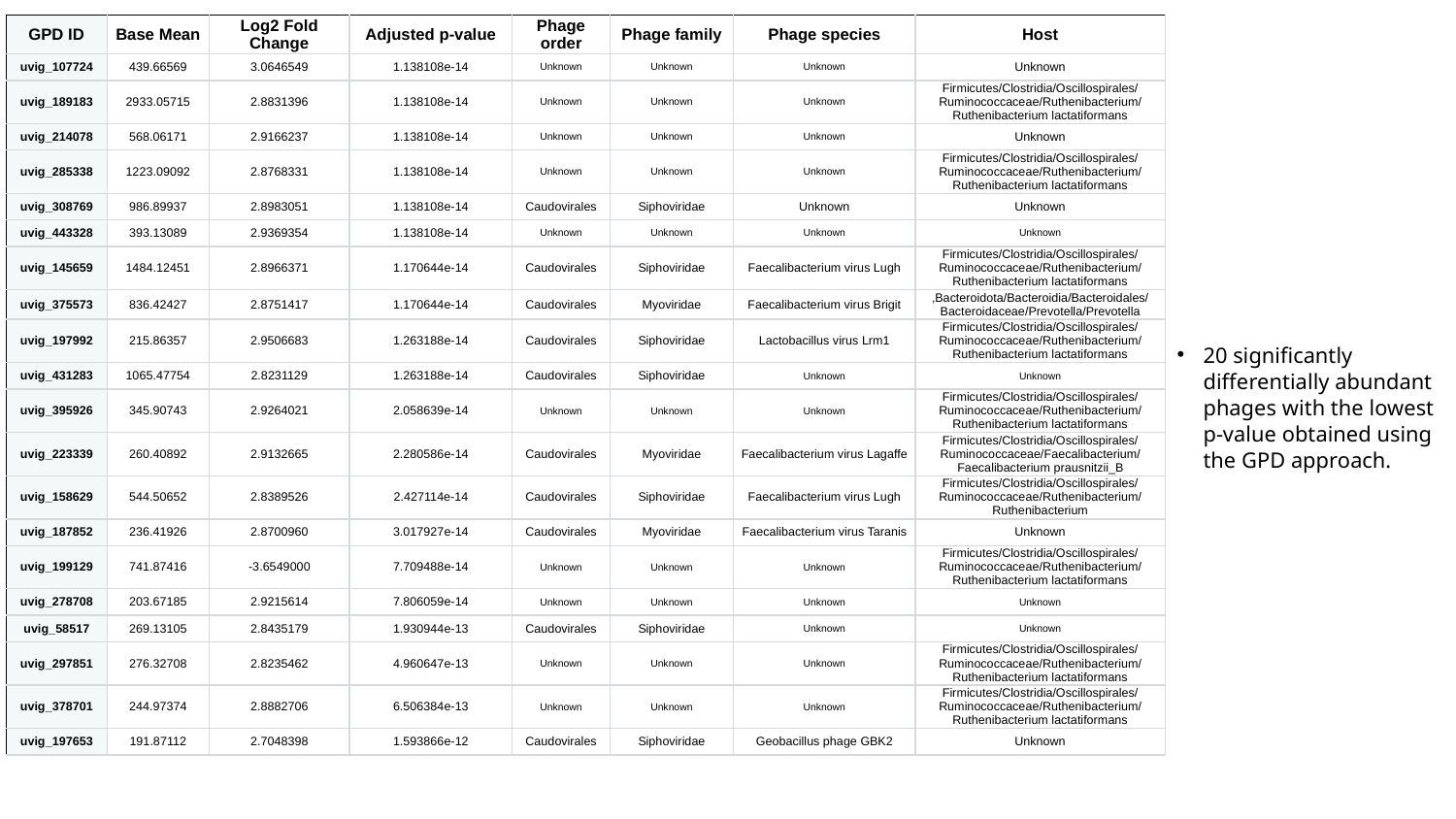

| GPD ID | Base Mean | Log2 Fold Change | Adjusted p-value | Phage order | Phage family | Phage species | Host |
| --- | --- | --- | --- | --- | --- | --- | --- |
| uvig\_107724 | 439.66569 | 3.0646549 | 1.138108e-14 | Unknown | Unknown | Unknown | Unknown |
| uvig\_189183 | 2933.05715 | 2.8831396 | 1.138108e-14 | Unknown | Unknown | Unknown | Firmicutes/Clostridia/Oscillospirales/Ruminococcaceae/Ruthenibacterium/Ruthenibacterium lactatiformans |
| uvig\_214078 | 568.06171 | 2.9166237 | 1.138108e-14 | Unknown | Unknown | Unknown | Unknown |
| uvig\_285338 | 1223.09092 | 2.8768331 | 1.138108e-14 | Unknown | Unknown | Unknown | Firmicutes/Clostridia/Oscillospirales/Ruminococcaceae/Ruthenibacterium/Ruthenibacterium lactatiformans |
| uvig\_308769 | 986.89937 | 2.8983051 | 1.138108e-14 | Caudovirales | Siphoviridae | Unknown | Unknown |
| uvig\_443328 | 393.13089 | 2.9369354 | 1.138108e-14 | Unknown | Unknown | Unknown | Unknown |
| uvig\_145659 | 1484.12451 | 2.8966371 | 1.170644e-14 | Caudovirales | Siphoviridae | Faecalibacterium virus Lugh | Firmicutes/Clostridia/Oscillospirales/Ruminococcaceae/Ruthenibacterium/Ruthenibacterium lactatiformans |
| uvig\_375573 | 836.42427 | 2.8751417 | 1.170644e-14 | Caudovirales | Myoviridae | Faecalibacterium virus Brigit | ,Bacteroidota/Bacteroidia/Bacteroidales/Bacteroidaceae/Prevotella/Prevotella |
| uvig\_197992 | 215.86357 | 2.9506683 | 1.263188e-14 | Caudovirales | Siphoviridae | Lactobacillus virus Lrm1 | Firmicutes/Clostridia/Oscillospirales/Ruminococcaceae/Ruthenibacterium/Ruthenibacterium lactatiformans |
| uvig\_431283 | 1065.47754 | 2.8231129 | 1.263188e-14 | Caudovirales | Siphoviridae | Unknown | Unknown |
| uvig\_395926 | 345.90743 | 2.9264021 | 2.058639e-14 | Unknown | Unknown | Unknown | Firmicutes/Clostridia/Oscillospirales/Ruminococcaceae/Ruthenibacterium/Ruthenibacterium lactatiformans |
| uvig\_223339 | 260.40892 | 2.9132665 | 2.280586e-14 | Caudovirales | Myoviridae | Faecalibacterium virus Lagaffe | Firmicutes/Clostridia/Oscillospirales/Ruminococcaceae/Faecalibacterium/Faecalibacterium prausnitzii\_B |
| uvig\_158629 | 544.50652 | 2.8389526 | 2.427114e-14 | Caudovirales | Siphoviridae | Faecalibacterium virus Lugh | Firmicutes/Clostridia/Oscillospirales/Ruminococcaceae/Ruthenibacterium/Ruthenibacterium |
| uvig\_187852 | 236.41926 | 2.8700960 | 3.017927e-14 | Caudovirales | Myoviridae | Faecalibacterium virus Taranis | Unknown |
| uvig\_199129 | 741.87416 | -3.6549000 | 7.709488e-14 | Unknown | Unknown | Unknown | Firmicutes/Clostridia/Oscillospirales/Ruminococcaceae/Ruthenibacterium/Ruthenibacterium lactatiformans |
| uvig\_278708 | 203.67185 | 2.9215614 | 7.806059e-14 | Unknown | Unknown | Unknown | Unknown |
| uvig\_58517 | 269.13105 | 2.8435179 | 1.930944e-13 | Caudovirales | Siphoviridae | Unknown | Unknown |
| uvig\_297851 | 276.32708 | 2.8235462 | 4.960647e-13 | Unknown | Unknown | Unknown | Firmicutes/Clostridia/Oscillospirales/Ruminococcaceae/Ruthenibacterium/Ruthenibacterium lactatiformans |
| uvig\_378701 | 244.97374 | 2.8882706 | 6.506384e-13 | Unknown | Unknown | Unknown | Firmicutes/Clostridia/Oscillospirales/Ruminococcaceae/Ruthenibacterium/Ruthenibacterium lactatiformans |
| uvig\_197653 | 191.87112 | 2.7048398 | 1.593866e-12 | Caudovirales | Siphoviridae | Geobacillus phage GBK2 | Unknown |
20 significantly differentially abundant phages with the lowest p-value obtained using the GPD approach.
