## Supplementary Table S13 for "Comparison of phage and plasmid populations present in the gut microbiota of Parkinson’s disease patients": Supplementary Table S13 - GPD base mean.pptx

### Slide 1
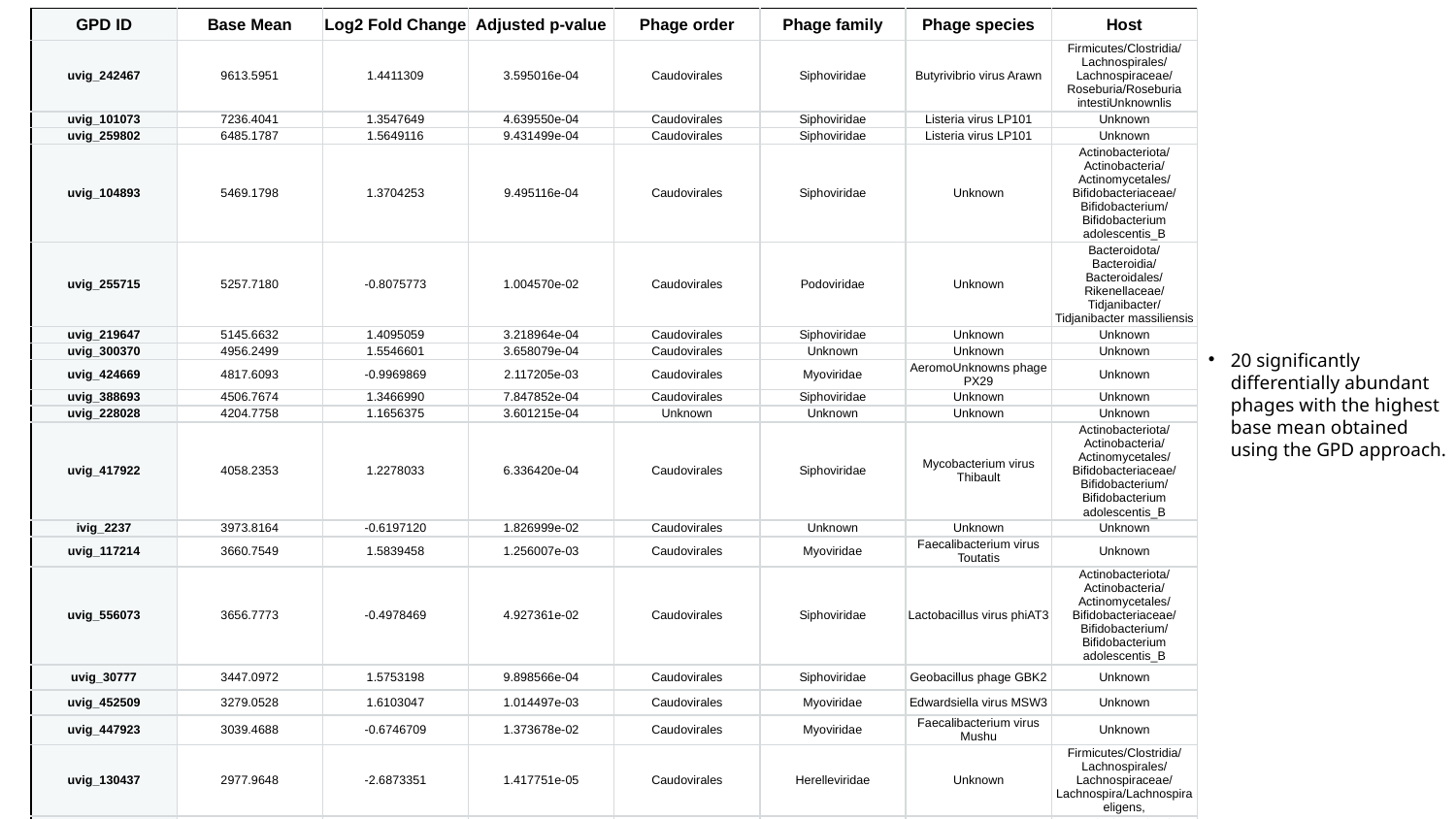

| GPD ID | Base Mean | Log2 Fold Change | Adjusted p-value | Phage order | Phage family | Phage species | Host |
| --- | --- | --- | --- | --- | --- | --- | --- |
| uvig\_242467 | 9613.5951 | 1.4411309 | 3.595016e-04 | Caudovirales | Siphoviridae | Butyrivibrio virus Arawn | Firmicutes/Clostridia/Lachnospirales/Lachnospiraceae/Roseburia/Roseburia intestiUnknownlis |
| uvig\_101073 | 7236.4041 | 1.3547649 | 4.639550e-04 | Caudovirales | Siphoviridae | Listeria virus LP101 | Unknown |
| uvig\_259802 | 6485.1787 | 1.5649116 | 9.431499e-04 | Caudovirales | Siphoviridae | Listeria virus LP101 | Unknown |
| uvig\_104893 | 5469.1798 | 1.3704253 | 9.495116e-04 | Caudovirales | Siphoviridae | Unknown | Actinobacteriota/Actinobacteria/Actinomycetales/Bifidobacteriaceae/Bifidobacterium/Bifidobacterium adolescentis\_B |
| uvig\_255715 | 5257.7180 | -0.8075773 | 1.004570e-02 | Caudovirales | Podoviridae | Unknown | Bacteroidota/Bacteroidia/Bacteroidales/Rikenellaceae/Tidjanibacter/Tidjanibacter massiliensis |
| uvig\_219647 | 5145.6632 | 1.4095059 | 3.218964e-04 | Caudovirales | Siphoviridae | Unknown | Unknown |
| uvig\_300370 | 4956.2499 | 1.5546601 | 3.658079e-04 | Caudovirales | Unknown | Unknown | Unknown |
| uvig\_424669 | 4817.6093 | -0.9969869 | 2.117205e-03 | Caudovirales | Myoviridae | AeromoUnknowns phage PX29 | Unknown |
| uvig\_388693 | 4506.7674 | 1.3466990 | 7.847852e-04 | Caudovirales | Siphoviridae | Unknown | Unknown |
| uvig\_228028 | 4204.7758 | 1.1656375 | 3.601215e-04 | Unknown | Unknown | Unknown | Unknown |
| uvig\_417922 | 4058.2353 | 1.2278033 | 6.336420e-04 | Caudovirales | Siphoviridae | Mycobacterium virus Thibault | Actinobacteriota/Actinobacteria/Actinomycetales/Bifidobacteriaceae/Bifidobacterium/Bifidobacterium adolescentis\_B |
| ivig\_2237 | 3973.8164 | -0.6197120 | 1.826999e-02 | Caudovirales | Unknown | Unknown | Unknown |
| uvig\_117214 | 3660.7549 | 1.5839458 | 1.256007e-03 | Caudovirales | Myoviridae | Faecalibacterium virus Toutatis | Unknown |
| uvig\_556073 | 3656.7773 | -0.4978469 | 4.927361e-02 | Caudovirales | Siphoviridae | Lactobacillus virus phiAT3 | Actinobacteriota/Actinobacteria/Actinomycetales/Bifidobacteriaceae/Bifidobacterium/Bifidobacterium adolescentis\_B |
| uvig\_30777 | 3447.0972 | 1.5753198 | 9.898566e-04 | Caudovirales | Siphoviridae | Geobacillus phage GBK2 | Unknown |
| uvig\_452509 | 3279.0528 | 1.6103047 | 1.014497e-03 | Caudovirales | Myoviridae | Edwardsiella virus MSW3 | Unknown |
| uvig\_447923 | 3039.4688 | -0.6746709 | 1.373678e-02 | Caudovirales | Myoviridae | Faecalibacterium virus Mushu | Unknown |
| uvig\_130437 | 2977.9648 | -2.6873351 | 1.417751e-05 | Caudovirales | Herelleviridae | Unknown | Firmicutes/Clostridia/Lachnospirales/Lachnospiraceae/Lachnospira/Lachnospira eligens, |
| uvig\_189183 | 2933.0571 | 2.8831396 | 1.138108e-14 | Caudovirales | Siphoviridae | Gordonia phage Schmidt | Actinobacteriota/Actinobacteria/Actinomycetales/Bifidobacteriaceae/Bifidobacterium/Bifidobacterium adolescentis\_B |
| uvig\_281278 | 2873.9822 | 1.4517249 | 5.840095e-04 | Caudovirales | Myoviridae | Faecalibacterium virus Mushu | Unknown |
20 significantly differentially abundant phages with the highest base mean obtained using the GPD approach.
