## Supplementary Fig. S11 for "Comparison of phage and plasmid populations present in the gut microbiota of Parkinson’s disease patients": Supplementary Fig. 11 - CRISPR .pptx

### Slide 1
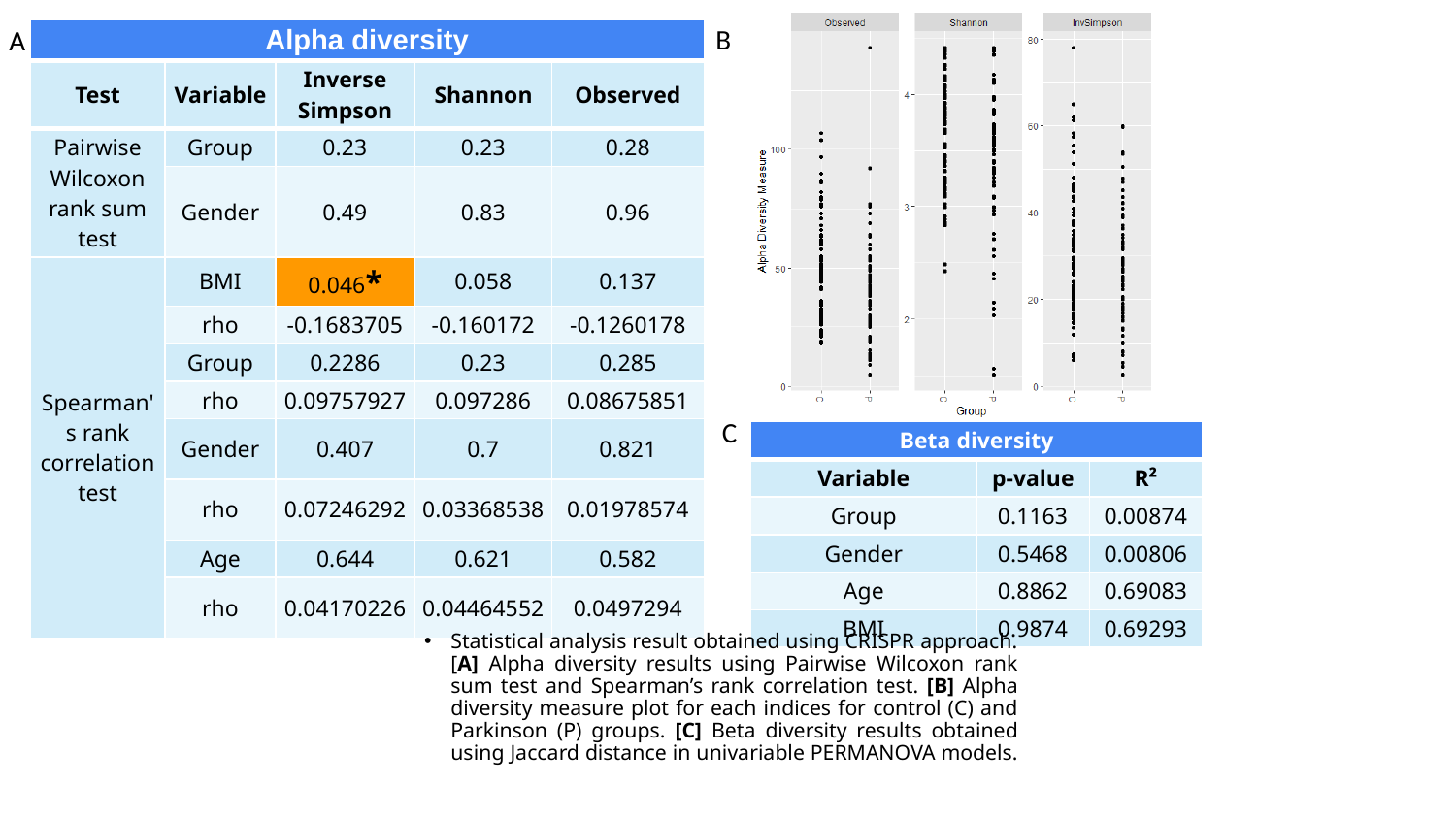

B
A
| Alpha diversity | | | | |
| --- | --- | --- | --- | --- |
| Test | Variable | Inverse Simpson | Shannon | Observed |
| Pairwise Wilcoxon rank sum test | Group | 0.23 | 0.23 | 0.28 |
| | Gender | 0.49 | 0.83 | 0.96 |
| Spearman's rank correlation test | BMI | 0.046\* | 0.058 | 0.137 |
| | rho | -0.1683705 | -0.160172 | -0.1260178 |
| | Group | 0.2286 | 0.23 | 0.285 |
| | rho | 0.09757927 | 0.097286 | 0.08675851 |
| | Gender | 0.407 | 0.7 | 0.821 |
| | rho | 0.07246292 | 0.03368538 | 0.01978574 |
| | Age | 0.644 | 0.621 | 0.582 |
| | rho | 0.04170226 | 0.04464552 | 0.0497294 |
C
| Beta diversity | | |
| --- | --- | --- |
| Variable | p-value | R² |
| Group | 0.1163 | 0.00874 |
| Gender | 0.5468 | 0.00806 |
| Age | 0.8862 | 0.69083 |
| BMI | 0.9874 | 0.69293 |
Statistical analysis result obtained using CRISPR approach. [A] Alpha diversity results using Pairwise Wilcoxon rank sum test and Spearman’s rank correlation test. [B] Alpha diversity measure plot for each indices for control (C) and Parkinson (P) groups. [C] Beta diversity results obtained using Jaccard distance in univariable PERMANOVA models.
